# Decoding gene regulation in the fly brain

**DOI:** 10.1101/2021.08.11.454937

**Authors:** Jasper Janssens, Sara Aibar, Ibrahim Ihsan Taskiran, Joy N. Ismail, Katina I. Spanier, Carmen Bravo González-Blas, Xiao Jiang Quan, Dafni Papasokrati, Gert Hulselmans, Samira Makhzami, Maxime De Waegeneer, Valerie Christiaens, Stein Aerts

## Abstract

The *Drosophila* brain is a work horse in neuroscience. Single-cell transcriptome analysis ^1–5,^ 3D morphological classification ^6^, and detailed EM mapping of the connectome ^7–10^ have revealed an immense diversity of neuronal and glial cell types that underlie the wide array of functional and behavioral traits in the fruit fly. The identities of these cell types are controlled by – still unknown – gene regulatory networks (GRNs), involving combinations of transcription factors that bind to genomic enhancers to regulate their target genes. To characterize the GRN for each cell type in the *Drosophila* brain, we profiled chromatin accessibility of 240,919 single cells spanning nine developmental timepoints, and integrated this data with single-cell transcriptomes. We identify more than 95,000 regulatory regions that are used in different neuronal cell types, of which around 70,000 are linked to specific developmental trajectories, involving neurogenesis, reprogramming and maturation. For 40 cell types, their uniquely accessible regions could be associated with their expressed transcription factors and downstream target genes, through a combination of motif discovery, network inference techniques, and deep learning. We illustrate how these “enhancer-GRNs” can be used to reveal enhancer architectures leading to a better understanding of neuronal regulatory diversity. Finally, our atlas of regulatory elements can be used to design genetic driver lines for specific cell types at specific timepoints, facilitating the characterization of brain cell types and the manipulation of brain function.

## Main

The brain consists of a myriad of different neuronal and glial types, each unique in their morphology and function. The *Drosophila* brain, which contains around 100,000 cells, is uniquely positioned as a model in which the diversity of brain cell types can be investigated. Recent advances in electron microscopy have allowed the creation of connectome maps of the different regions in the *Drosophila* brain ^7–10^ , while the availability of genetic driver lines ^11^ provides genetic access to many cell types for understanding neuronal function ^12^. Furthermore, this diversity of cell types has been bolstered by single-cell transcriptomics on the adult brain ^1–5^, the larval brain ^13–15^, and the ventral nerve cord ^16^. The recent development of single-cell assay for transposase accessible chromatin by sequencing (scATAC-seq), makes it possible to measure chromatin accessibility of single cells in high throughput ^17, 18^, providing an additional crucial layer of information underlying neuronal identity: which genomic regions encode the regulatory information to create and maintain each cell type. The integrated analysis of transcriptomics and chromatin accessibility makes it then possible to jointly study enhancers and gene expression to discover precise regulatory programs across cell types ^19–21^.

Cell type identity is defined by the activity of GRNs in which combinations of transcription factors activate or repress target genes. In Davie et al., we showed that for many different cell types in the brain, unique transcription factor (TF) combinations can be identified that govern gene regulatory networks, thus highlighting the unique role TFs play in neuronal fate determination. Neural progenitors have been shown to exist along two axes of differentiation, one temporal and one spatial, and both are guided by transcription factor changes ^22, 23^. This patterning of neural progenitors is presumed to give rise to the diverse cell types found in the adult brain. The combinatorial expression of transcription factors also governs key neuronal features by guiding dendritic targeting and neurotransmitter determination ^2, 24–27^ and changing expression of a single TF can lead to a change in neuronal fate ^28, 29^. Whereas transcriptomic studies in *Drosophila* have led to the inference of TFs and their putative target genes, they suffer from high false positive rates. Moreover, TF activity often cannot be predicted from the transcription level of TFs, as it depends on a large number of variables ^30^ such as protein activity, protein localization, and the presence of co-binding TFs and co-factors. On the other hand, profiling chromatin accessibility leads to a direct read-out of possible TF binding sites and therefore bypasses these limitations ^31^.

Here, we build a single-cell multi-omics atlas across the development of the fly brain, including transcriptomic and chromatin accessibility profiles from late larva to adult, covering the dynamic processes of neurogenesis, maturation, and maintenance. We identify key regulators of neuronal and glial cell identity, decipher the enhancer code for specific neuronal subtypes, and generate “informed” enhancer driver lines that allow cell type specific manipulation. This information is available as a resource at http://flybrain.aertslab.org to allow users to explore the data in detail for their own cell populations of interest.

### Unique chromatin landscapes underlie neuronal diversity

To study the regulatory programs of neuronal diversity, we profiled chromatin accessibility of 240,919 cells from the entire brain using single-cell ATAC-seq (10x Chromium). We dissected brains at nine time points from third instar larvae to young adult flies, covering the most important stages of neuronal development ^26, 32^ (Fig. 1a). The experiments were carried out with four different wild type polymorphic strains ^33^, allowing a higher number of nuclei per run, while detecting and removing doublets (see Methods, and Extended Data Table 1). In addition, to enrich for central brain cell types that are often hard to detect ^1, 3^, we performed two additional runs on adult brains without the optic lobes as these contain more than two thirds of all brain cells in numbers, but with more reduced diversity than the central brain. We complement this chromatin atlas with previously published scRNA-seq data from the larval brain (5,054 cells) ^15^ and we updated our scRNA-seq atlas of the adult brain by expanding our previous atlas of 56,902 cells ^1^ to 118,687 high quality cells. In this enlarged scRNA-seq dataset, we identified 204 clusters of which 66 could be annotated as known cell types (see Methods).

**Figure 1:**
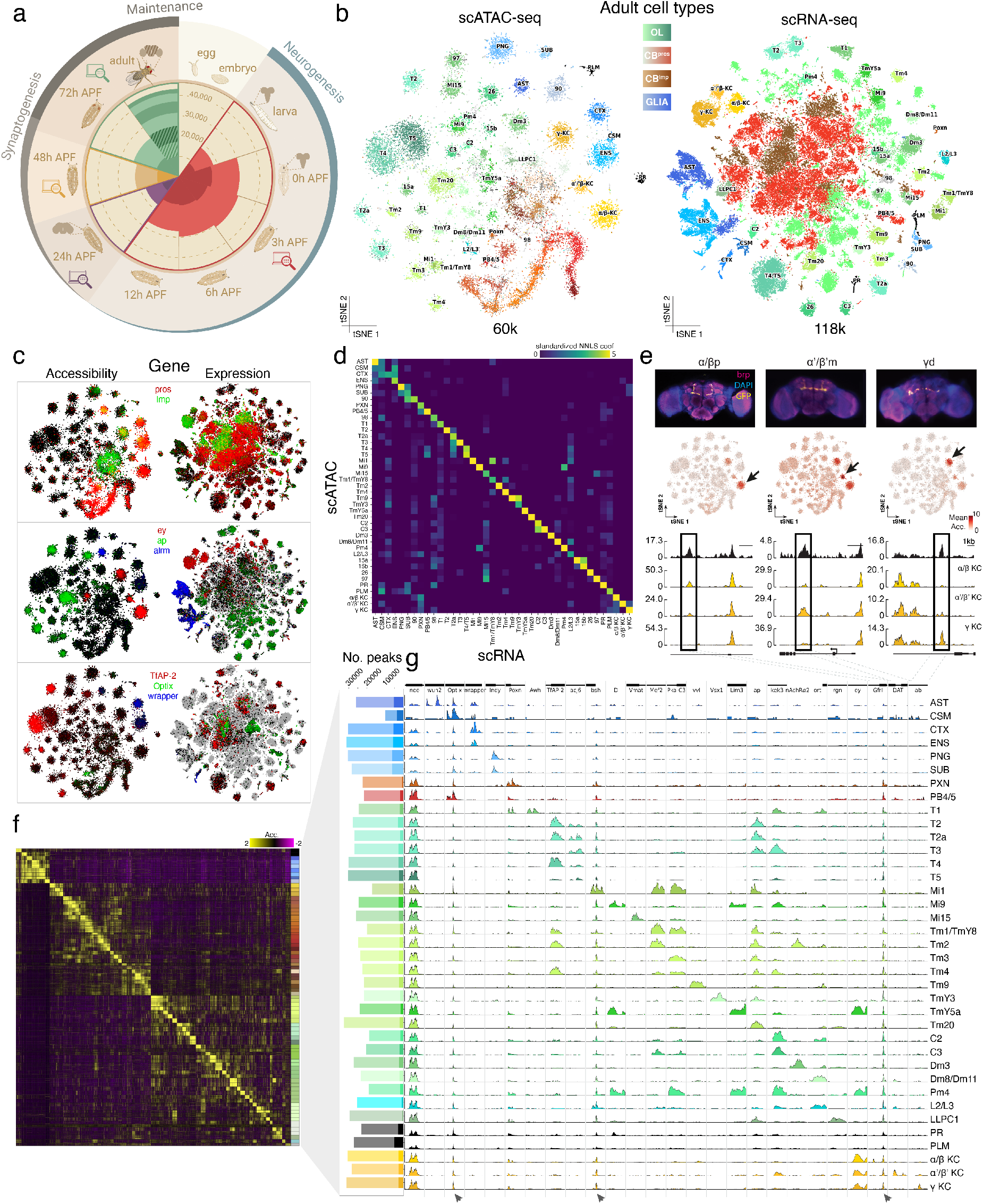
Overview of the chromatin atlas and characterization of the chromatin landscape of the adult cell types. **a.** Experimental overview showing number of cells per timepoint, covering the spectrum of neurogenesis and synaptogenesis. Runs are demarcated by color switches; central brain only runs are accentuated. **b.** 2D projections (tSNE) of the adult cell types in scATAC-seq (left, includes adult and 72h APF cells) and scRNA-seq (right, adult cells only). **c.** Gene accessibility and gene expression display a similar pattern for many genes. **d.** Heatmap of NNLS regression coefficients after z-normalization showing the correspondence between RNA and ATAC clusters. **e.** Overview of bulk ATAC-seq on three sorted cell populations. (top) Confocal images of three Kenyon cell subtypes targeted with split-GAL4 lines; (middle) tSNE showing accessibility of cell type specific regions; (bottom) profiles show similarity between bulk ATAC-seq (black) and scATAC-seq. **f.** Overview of differentially accessible regions per cluster, bar plot shows number of accessible regions per cell type with cell type specific regions in dark. **g**. Aggregate profiles of differentially accessible regions (max value=50 (30 for *rgn* and *ab*)). Note the constitutively accessible regions in *Optix*, *bsh* and *Gfrl* (arrows). Cell type abbreviations: Astrocyte-like glia (AST), Chiasm glia (CSM), Cortex glia (CTX), Ensheathing glia (ENG), Perineurial glia (PNG), Sub-perineurial glia (SUB), Poxn-neurons of the ellipsoid body (PXN), Protocerebral bridge neurons (PB), Photoreceptors (PR), Plasmatocytes (PLM), Kenyon Cells (KC). Numbers are unidentified clusters that match between scRNA- and scATAC-seq.

We first analyzed the open chromatin landscape in the adult cell types, as they represent the tips of the developmental manifold. To this end, we combined the 60,624 cells from adult and late stage pupa (72h APF), as these stages are very similar and most neurogenesis and circuit assembly has finished (Extended Data Fig. 1). The analysis with cisTopic ^34^ resulted in four main categories of cells (Fig. 1b, glia, optic lobe, Kenyon cells, and the remainder of the central brain) that we further subclustered in 79 clusters (Extended Data Fig. 2, and Methods). To link these clusters to specific cell types, we exploited the expanded single-cell transcriptome atlas of the adult brain. We generated a gene accessibility matrix by taking the sum of the accessibility of the regions within the gene body and upstream of its TSS, weighted by distance and variability (Fig. 1c, Extended Data Fig. 3a). Co-clustering the RNA and gene accessibility datasets allowed us to annotate 35 clusters (Extended Data Fig. 3b-d). To match and confirm additional clusters across modalities, we used marker gene enrichment ^35^ and non-negative least squares (NNLS) regression ^36^ (Fig. 1d, Extended Data Fig. 3e). The combination of these methods led to a final annotation of 43 of the 79 clusters, which were unambiguously one-to-one linked to RNA.

The annotated cell types include six glial subtypes (∼10-15%, in blue in Fig. 1b) as well as non-brain cells (∼1%, plasmatocytes and photoreceptors, in black); but, as expected, the majority of cells are neurons (∼85-90%; optic lobe in green, central brain clusters in red). Interestingly, optic lobe neurons form clear and distinct clusters, while central brain neurons appear as a continuum. This may be explained by the organization of the optic lobe, in which cell types are present in multiple copies, forming repetitive columnar structures that process input from the 800 ommatidia of the compound eye. While many unicolumnar cell types (present in every column) were identified, we were unable to annotate the sparser multicolumnar neurons (connecting multiple columns). In the central brain, we identified the three Kenyon cell (KC) subtypes and two smaller cell types of the central complex: ring neurons of the ellipsoid body, and the protocerebral bridge. This is similar to observations in transcriptome data of the central brain, where central brain cell types and multicolumnar optic lobe neurons were more difficult to identify due to their low cell numbers ^1, 3^. Intriguingly, central brain clusters are split into *Imp*^+^ or *prospero* (*pros*)^+^ cells based on the scATAC-seq based gene accessibility, recapitulating differences shown previously by scRNA-seq in the brain ^1^, ventral nerve cord ^16^, and larval brain ^15^ (see also further below).

To further validate cell type annotation and their associated regulatory regions, we used split-GAL4 lines to label the three KC subtypes ^37^ and performed bulk ATAC-seq after FAC-sorting ^38^ (Fig. 1e). The differential regions from the bulk profiles matched the single-cell clusters (Fig. 1e: cell type specific accessible regions highlight matching clusters in the tSNE). Comparing the scATAC-seq aggregated profile with the sorted bulk data profiles confirms the high concordance at marker regions, as illustrated in Fig. 1e, with the aggregates of αβ-KCs having matching peaks near *Gfrl*, α’β’-KCs near *DAT*, and γ-KCs near *ab*, thus confirming the high quality of cell type specific chromatin accessibility based on scATAC-seq.

Although not all clusters were mapped to specific cell types, each cluster has a unique chromatin accessibility profile, with a range of 105 to 4,732 differentially accessible regions (DARs) out of a total of 24,543 median accessible regions per cluster or cell type (Fig. 1f-g). Accessibility of intronic and distal intergenic regions correlated better with gene expression compared to the accessibility of the promoter/TSS region, confirming previous observations ^39^(Extended Data Fig. 4). Indeed, many broadly used and validated marker genes have their TSS ubiquitously accessible and their specificity may be controlled by more distal regions (e.g., *bsh* in Mi1 neurons, *Optix* in PB neurons [arrows in Fig. 1g]). Of the 1,017 DARs of the mushroom body Kenyon cells that overlap with tested enhancer-reporter regions in the Janelia FlyLight collection, 588 are reported as active enhancers in the mushroom body (Extended Data Fig. 5) suggesting that DARs often function as enhancers.

Interestingly, while T4 and T5 neurons are grouped into the same cluster in the transcriptome data their chromatin profiles split them into two distinct clusters, even though there are only 110 differentially accessible regions between T4 and T5. Three of the regions accessible in T4 neurons are located in the locus of *TfAP-2*, a transcription factor that is specifically expressed in T4 ^40^. Further sub- clustering reveals that a/b and c/d subtypes can also be separated by chromatin accessibility (Extended Data Fig. 6). Given this high resolution in the scATAC-seq data, it is surprising that 60K cells were not sufficient for identification of all cell types in the central brain. For example, the olfactory projection neurons (OPNs) form a separate cluster in the 57K scRNA-seq dataset ^1^ but not in the 60K scATAC-seq dataset. Further increasing the cell count to 88.3K cells by including the 48h time point, where OPNs are already present, does not resolve this issue (Extended Data Fig. 2e). OPNs and other central brain cell types are thus harder to distinguish at the level of accessible chromatin, compared to the level of the transcriptome.

In conclusion, scATAC-seq of the fly brain yields high-quality cell type specific chromatin accessibility profiles that correspond to matching transcriptomes, allowing all large cell types in the optic lobes and multiple cell types in the central brain, including the three KC subtypes, to be characterized by an associated set of differentially accessible regions.

### Concordant TF expression, enhancer accessibility, and gene expression yield enhancer- GRNs

Current descriptions of gene regulatory networks have been mostly focused on imputing transcription factors with their target genes by co-expression, sometimes enhanced with motif detection ^35, 41–45^ The availability of transcriptome *and* chromatin accessibility profiles of the matched cell types opens up the possibility of exploring their regulatory code in much greater detail. In particular, we aim to unravel high-confidence cell type specific “enhancer-GRNs” (eGRNs), including the key transcription factors of a cell type, as well as their target genes and the enhancers through which they are regulated.

To reconstruct eGRNs, we developed a computational strategy that exploits the matched scRNA- and scATAC-seq data to identify candidate transcription factors that are both expressed and have their recognition motif enriched in the open regions of a given cell type (Fig. 2a). As a first step, we defined “cistromes” (i.e., candidate target regions) for each transcription factor: the subset of DARs of each cell type, in which the TF is expressed and its motif is significantly enriched (Fig. 2b-c) (see Methods). We found cistromes for 206 TFs (out of the 251 expressed TFs with a motif in our database), including TFs with pan-neuronal, pan-glial, and cell type-specific activity (89 of them shown in Fig. 2b).

**Figure 2:**
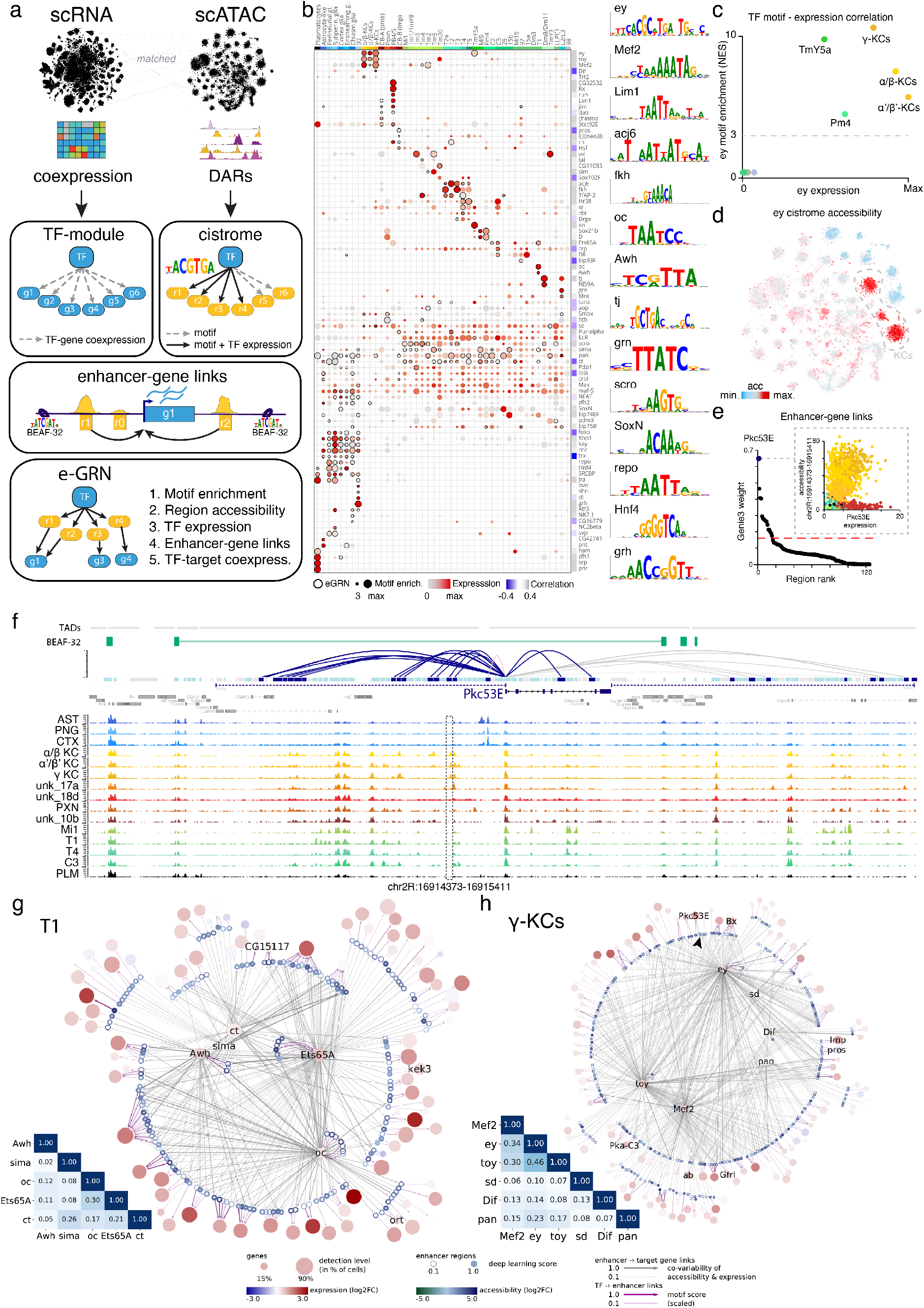
Construction of eGRNs for key brain cell types through multi-omic data integration. **a.** A multi-omic approach for the creation of e(nhancer-)GRNs. **b.** Dotplot showing expression and motif enrichment for the TFs within an eGRN (marked circles show the presence of a cell type-specific eGRN). **c.** Ey motif enrichment versus expression, per cell type. **d.** tSNE showing the accessibility of the regions of the Ey-cistrome across cell types in the adult. Note high accessibility in the *ey*-expressing Kenyon cells. **e.** Regulatory region selection for *Pkc53E*; the inset shows the accessibility of the top region versus the gene expression (input to the random-forest), regions with a weight above the threshold are linked to the gene. **f.** Overview of selected tracks in the *Pkc53E* locus; only the enhancer-gene links between two BEAF-32 peaks (green bar) are kept. **g-h.** eGRNs for T1 neurons and γ -KCs (regions are coloured in blue shades, genes in red; regulatory TFs are in the center). Insert heatmaps show Jaccard index between TF target regions. Arrow points to *Pkc53E* enhancer shown in **f**.

The full list of predicted regulators and their cistromes can be downloaded from the web portal (https://flybrain.aertslab.org). These include key regulators for KCs, in which we confirmed Mef2 for γ-KCs and αβ-KCs ^46, 47^, in addition to Ey (Fig. 2d). For ellipsoid body neurons, Grain (grn) and Dichaete (D) emerged as key regulators ^48^. In T1 neurons, we identified Ets65A ^40^, and Ocelliless (oc) as main regulators ^49^ and combinations of Acj6, Fkh, TfAP-2, and SoxN/Sox102F in the other T-neurons ^2, 4, 27, 40^. Glial cells show *repo* and *Kay* expression, and their respective motifs are enriched in glial DARs ^50^. Interestingly, we also identify TFs with a negative correlation between gene expression and motif enrichment, suggesting a repressive role for instance for Pros, Lola and Cut (ct) (Fig 2b).

In the second step, we linked the cistrome regions to their target genes. We approached this by calculating a co-variability score for the *expression* of each gene and the *accessibility* of the regulatory regions nearby (using random-forest regression, and “metacells” to match the cell types across data modalities, see methods). Previous work has shown that regulatory interactions can occur over large distances but are mostly confined within chromatin domains, in so-called “genomic regulatory blocks” (127kb median size, ^51^), a HiC-derived topological associated domain (TAD, 13kb median size, ^52^), or between two BEAF-32 boundary elements (57kb median distance, ^53^). Taking this into account, we considered enhancer-gene interactions in a window of >100kbp around each gene (50kbp up- and down-stream, plus introns) (Fig. 2e), which led to an average of 9 linked regions per gene (Fig. 2f), 55% of which lie between BEAF-32 boundaries (Extended Data Fig. 7).

These enhancer-gene links provide a set of potential target genes per cistrome. However, to reduce the rate of false positives and obtain a higher-confidence network, as a third and final step, we tested whether the expression of each set of target genes co-varies with the predicted TF, similar to the principles of SCENIC ^35^ (based on gradient boosted regression and gene set enrichment analysis, see Methods). This confirmed that when the analysis is restricted to links that are within BEAF-32 domains, the correlation with TF expression is of slightly higher quality, allowing us to select the 89 highest-confidence cistromes (highlighted by a dark border in Fig. 2b) together with their top ranked target genes, which form the “enhancer-GRN” for the different cell types (Fig. 2g).

The eGRNs for the 40 available cell types have an average of 6 TFs, collectively regulating 108 target genes through 138 enhancers within BEAF-32 boundaries. Fig. 2g shows the network of T1 neurons, which highlights the combination of 5 TFs (Ets65A, Oc, Ct, Sima and Awh) regulating between 15 and 60 targets, with Oc and Ets65A auto-regulating, and around 50% of the targets being co-regulated by at least two TFs. Similarly, the network for γ-KCs (Fig. 2h) reveals Mef2 and Ey/Toy as auto-regulatory key factors that regulate 77 to 83 genes, alongside new candidate TFs Dif, Sd, and Pan regulating an average of 29 genes. 2/3 of the KC target genes are co-regulated by a minimum of two TFs, specifically revealing a high overlap between Ey/Toy and Mef2. The analysis of the 40 eGRNs also suggests that, while at least 57% of the genes are regulated by several regions within the same cell type, 95% of the regulatory interactions involve multiple TF inputs that occur through a common enhancer. In conclusion, we have developed a new computational strategy and constructed 40 cell type specific “enhancer-GRNs” with key transcription factors, target genes and enhancers (Extended Data Fig. 8).

### Combinatorial TF expression is reflected by enhancer architecture

Enhancer-GRNs provide intuitive insight into the regulatory state of a cell type and highlight the combinatorial nature of TF-target interactions. To further investigate how TF ensembles yield highly accurate spatio-temporal gene expression patterns, we examined the sequence of the predicted enhancers in detail using a deep learning model and tested their activity by a selection of *in vivo* enhancer-reporter assays. We focused on a subset of the data that includes KCs, T-neurons, and glia and we re-analyzed these with cisTopic (Fig. 3a). cisTopic provides both a cell clustering and a region clustering in the form of topics. These topics are sets of regions that can be accessible in specific cell types or in multiple (Fig. 3b). We then trained a convolutional neural network using the sequences of the topics as input in order to predict in which cell types they are accessible ^54^ (see Methods, Fig. 3c, Extended Data Table 2). Evaluation of the model’s accuracy using cross-validation and left-out test data, yielded accurate classification of promoters, BEAF-32 boundary elements (3 topics, average auPR=0.36), pan-neuronal and pan-glial regions (5 topics with auPR=0.30 and 2 topics with auPR=0.30 respectively), and cell type specific enhancers for KCs, the glial subtypes, and each of the T-neuron classes (Fig. 3b, Extended Data Table 3).

**Figure 3:**
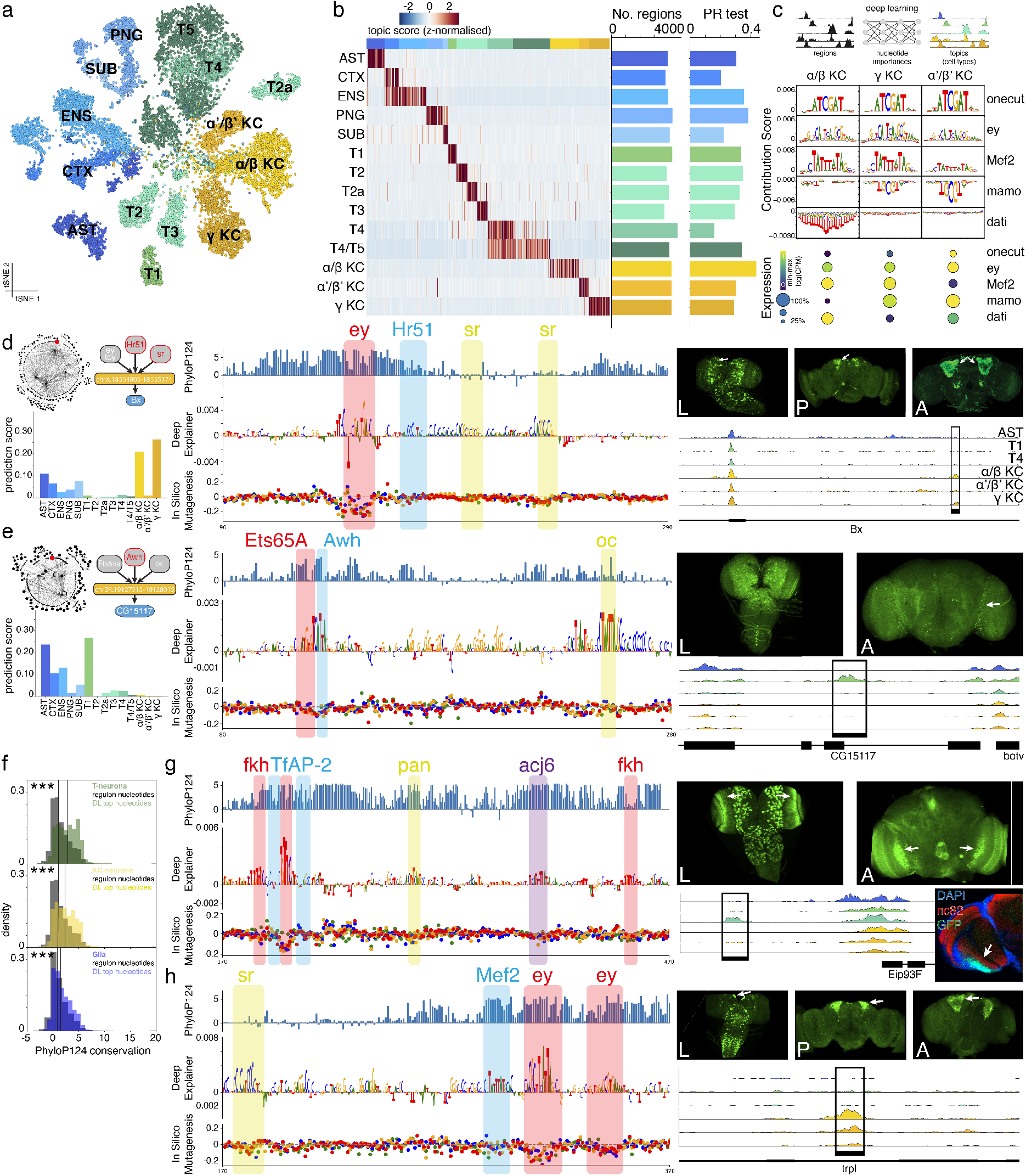
Deep learning unravels enhancer make-up. **a.** tSNE of cisTopic analysis on subset of Kenyon cells, T-neurons and glia. **b.** Topic heatmap showing cell type specific topics. Bar plots show number of regions per topic (cutoff p=0.995) and area under precision-recall curve (auPR) of deep learning model, validated on test dataset. **c.** Architecture of deep learning model in which regions are assigned to topics through nucleotide contributions. Contributions of the nucleotides lead to de-novo motif discovery, showing motifs of known KC factors, matching their expression. Note that negative nucleotide importance leads to repression of accessibility. **d.** Selected region from the γ-KC eGRN is accurately predicted to belong to a γ-KC topic. DeepExplainer of a subset of the region view highlights Ey, Hr51 and Sr binding sites. Cloning of the region (black box) leads to KC-specific expression patterns. **e.** Selected region from the T1 eGRN is accurately predicted to belong to a T1 topic. DeepExplainer of a subset of the region view highlights Ets65A, Awh and Oc binding sites. Cloning of the region leads to T1-specific expression patterns in the adult. **f.** Histograms showing the conservation of nucleotides in the T-neurons, KC and glia eGRN enhancers: selected by the DL model (colored, n= 12,429 (T), 11,625 (KC), 53,007 (Glia)) against the whole region (grey, n= 486,500 (T), 452,500 (KC), 2,125,500 (Glia)), lines showing the median. Contrast was performed with two-sided Wilcoxon rank-sums test (stars mark pval<1e-10). **g.** DAR in T4 neurons leads to the creation of a specific T4 driver line. DeepExplainer of a subset of the region view highlights Fkh, TfAP-2, Pan and Acj6 binding sites. Cloning of the region leads to T4-specific expression patterns. **h.** DAR in KCs leads to the creation of a specific KC driver line. DeepExplainer of a subset of the region view highlights Mef2, Sr and Ey binding sites. Cloning of the region leads to KC-specific expression patterns.

To reveal the TF motifs that underlie cell type-specific accessibility, DeepExplainer ^55^, which identifies the importance of each nucleotide on a given sequence for the final prediction, and TF-Modisco ^56^, which uses the nucleotide importance scores and identifies motifs from reoccurring sequence patterns with high nucleotide importance score, were used to derive key features per topic. KC regions are predicted using motifs of Ey, Onecut, Mef2, Mamo and Dati, matching their high gene expression levels in KC (Fig. 3c). Thus, the deep learning model adds additional TFs, enhancers, and target genes to the KC eGRN. Interestingly, two of these, namely Mamo and Dati, have negative nucleotide importances, meaning that the presence of their motif is correlated with a closing of the peak, which may reflect a repressive function. The most important motifs linked to candidate TFs for T-neurons include Fkh, TfAP-2, Acj6 and Ct, Repo, Zfh2 and Klu for glia (Extended Data Fig. 9).

The eGRN framework we presented above, complemented with the deep learning framework, led to the prediction of cell type specific enhancers. We selected 52 genomic regions (of which 27 are nodes in the eGRNs, Extended Data Table 4, Extended Data Fig. 10a-b) for which we created a transgenic GFP reporter line (see Methods). For instance, a region associated with the *Bx* gene in the γ-KCs eGRN (Fig. 3d) is accurately predicted by the deep learning model, which reveals a candidate Ey binding site and additional motifs that we could match with Hr51 and Sr binding sites. While these two factors are associated with KC cistromes, their target genes did not pass the eGRN association filters, but using deep learning we recovered these TFs. The reporter activity of the enhancer is detected mainly in KCs, matching with the cell type specificity of the peak. Nevertheless, there is some off-target GFP signal in small cell populations in the central brain that are not part of the current clusters due to the lower resolution of cell types there. Another cloned region is predicted to activate *CG15117* expression in T1 neurons (Fig. 3e). The model correctly predicts the region as a T1 enhancer, using Oc and Ets65A motifs, and predicts an additional Awh binding site. The reporter activity shows specific activity in T1 neurons, matching the specificity of *CG15117* expression. Interestingly, both examples display different expression patterns in the larval brain compared to the adult brain, suggesting a recycling of the same enhancer in different cell types through development. Inspecting the conservation of eGRN enhancers using whole-genome alignments across 124 different insect species ^57^, reveals that the informative nucleotides predicted by the model show a higher conservation in each of the three main classes (KC, glia and T neurons, Wilcoxon p<1e-10), with the highest conservation in T neurons, suggesting a functional importance of the predicted binding sites (Fig. 3f).

Based on these results, we envision a potential use for this atlas as the starting point for the design of reporter lines to target specific cell populations throughout development. Although we could partially validate this potential by taking advantage of Janelia’s FlyLight ^11^ and Vienna Drosophila Resource Center (VDRC) enhancer lines (Extended Data Fig. 5) the subsequent interpretation is compromised due to their length, since the Janelia and VDRC regions are 2-3kb long and often contain multiple predicted enhancers. Using our transgenic lines for 52 candidate enhancers (including the examples shown in Fig. 3), where only a 300-1732bp ATAC-seq peak was cloned, we could achieve higher specificity compared to these larger regions. Indeed, we confirmed that for Janelia regions that have only one ATAC-seq peak, the active enhancer resides within the ATAC peak, which recapitulates the correct pattern similar to the larger encompassing Janelia selection (Extended Data Fig. 11a-b). In addition, we found that Janelia lines with multiple ATAC-seq peaks show activity in multiple cell types, while the activity of the individually cloned peaks becomes specific (Extended Data Fig. 11c). Finally, we confirm that split-GAL4 lines, which combine two enhancers in an “AND” logic, can be recapitulated as the intersection of their ATAC-seq signal (Extended Data Fig. 11d).

Using complementary approaches of eGRN inference and deep learning is useful, since the eGRN is based on enrichment analyses, region-to-gene mappings, and thresholding, and is biased to TF-target activation. The deep learning model identifies candidate repressors such as Dati and Mamo, predicts additional DARs as enhancers (Fig. 3g-h), and highlights binding sites in T4 and KC enhancers.

Many of the cloned enhancers show activity in the larval brain, with some enhancers having a broader expression pattern, suggesting developmental roles for many adult enhancers (Extended Data Fig. 10c-d). Surprisingly, DARs and eGRN regions reach similar levels of activity, although correlation with gene expression is a major factor to determine activity (Extended Data Fig. 10d-f). This suggests that even though the eGRN annotation is a set of active enhancers where both transcription factor and target gene is inferred, many functional enhancers remain to be investigated. Overall our results yield similar rates compared to the embryo ^58^ and eye-antennal disc ^59^.

In conclusion, we have created a deep learning model that can unravel the logic behind the enhancers in eGRNs and in accessible regions. The binding sites predicted by the model match motifs found by conventional methods, and add additional ones, thereby increasing our recall. Finally, we show how this atlas can be used to create new driver lines starting from the eGRNs or from specifically accessible regions. Existing driver lines that are “noisy” can be separated into smaller functional components that can be used as standalone, more specific driver lines.

### Dynamic changes in chromatin accessibility during brain development

To investigate how adult neuronal diversity is generated, we studied dynamic chromatin accessibility changes during development. We first focused on neurogenesis by separately analyzing the timepoints ranging from third instar larval to 12h after puparium formation (APF), with a total of 135,275 cells. Note that for these time points, we dissected the entire central nervous system (including brain and ventral nerve cord). The analysis of these cells with cisTopic resulted in 200 topics and 54 cell clusters (Fig. 4a).

**Figure 4:**
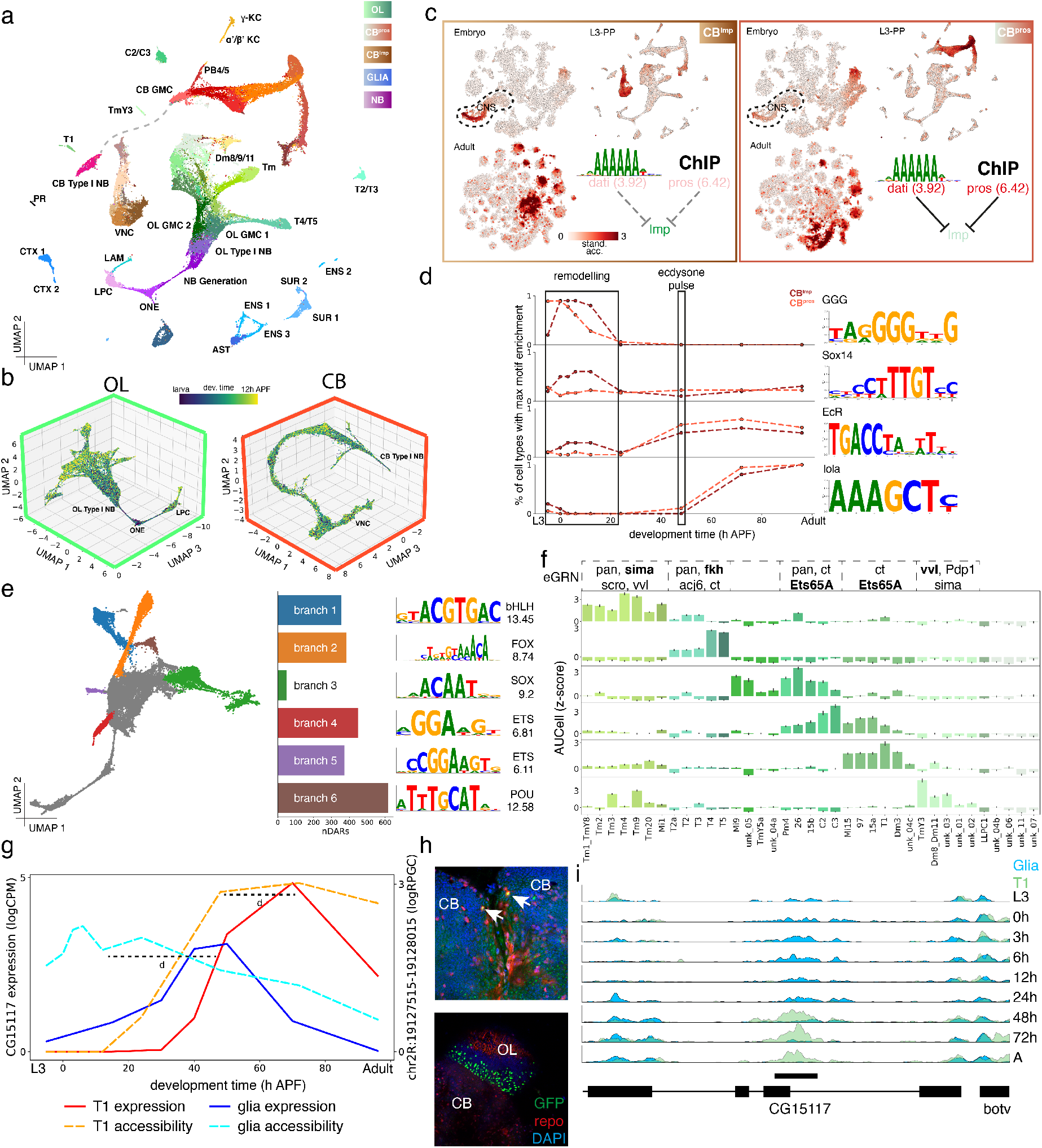
Development of neuronal cell types from neuroblasts through pupation. **a.** cisTopic UMAP of 92,954 cells for the analysis of early timepoints (third instar larvae to 12h APF, cells>900 FIP). Grey lines connect central brain neuroblasts to the central brain branch. **b.** Optic lobe and central brain branches in 3D-UMAP. **c.** Central brain and VNC duality between *pros* and *Imp*. **d.** Trajectory of motif enrichment in *Imp*^+^ and *pros*^+^ cells through development, showing modules for remodeling and an ecdysone response **e.** Six branches can be identified in the 3D optic lobe UMAP, each having their own unique accessibility regions. Motif enrichment reveals one transcription factor family per branch. **f.** Adult optic lobe cell types can be traced back to OL branches by enrichment of branch-specific regions through AUCell. TFs with eGRNs that overlap more than 10 regions with the branch are shown (matching motifs in bold). **g.** Example of an enhancer-switch between glia and T1 neurons, both in enhancer accessibility (full line) and target gene expression (dotted line). **h.** Staining of repo (red) and *CG15117* enhancer activity (GFP, green), showing switch from glia in early development (top, repo positive, central brain, 15h APF) to T1 neurons (bottom, repo negative, optic lobe, adult)). **i.** Track showing the dynamics of enhancer-switch between glia and T1 neurons. Cell type abbreviations: Astrocyte-like glia (AST), Cortex glia (CTX), Ensheathing glia (ENG), Surface glia (SUR), Ganglion mother cells (GMC), Lamina precursor cells (LPC), Lamina neurons (LAM), Optic lobe neuroepithelium (ONE), Neuroblasts (NB), Poxn-neurons of the ellipsoid body (PXN), Protocerebral bridge neurons (PB), Photoreceptors (PR), Plasmatocytes (PLM), Kenyon Cells (KC), Ventral nerve cord (VNC).

To annotate the cell clusters, we recursively trained an SVM classifier on the adult cell types and used it to label cell types in earlier stages, similar to recent work on RNA datasets ^4^ (Extended Data Fig. 12a-b). This data shows that adult DARs can be used to identify pupal cells, thus suggesting that neuronal fate is determined early on in development. To elaborate on this, we calculated core sets of regions per cell type that are continuously accessible at each timepoint, specifically characterizing cell types (Extended Data Fig. 12c-d). Similar to RNA-seq where in OPNs and optic lobe neurons, a maximum number of differentially expressed genes was detected in 48h APF and a minimum in the adult ^4, 32^, we find a decrease in DARs over time, and with a relative spike at 48h APF during synaptogenesis (Extended Data Fig. 12e).

On the other hand, the cell clusters that cannot be classified using the adult SVM are, as expected, the progenitor cell types. These clusters are characterized by accessible regions near the neuroblast markers deadpan (*dpn*) and asense (*ase*) (Extended Data Fig. 13a-b). During larval and pupal neurogenesis, the optic lobe neuroepithelium (ONE) in the outer proliferation center is converted to medulla neuroblasts (NB) and lamina precursor cells (LPCs), while quiescent embryonic central brain neuroblasts are reactivated ^60^. To reveal the dynamics of the bifurcation of neuroepithelium into either medulla NBs or LPCs, we fitted a trajectory through the UMAP and calculated modules for the different steps, calculating a complementary scRNA-seq trajectory from larval data ^15^ (Extended Data Fig. 13c-d). In the ONE state, grainyhead (*grh*) and zelda (*zld*) are found as major regulators, but are quickly replaced by glial cells missing (*gcm*) when progressing towards the LPC fate ^61^. The differentiation into medulla neuroblasts co-occurs with a phase with pointed (*pnt*) and scute (*sc*) ^60^, followed by *E(bx)* and finally the emergence of a neuro-GGG motif ^58, 59^ and known neuroblast factors *Ham* and *Kr* ^62, 63^. To study the differences between neural progenitors in more detail, we performed motif enrichment on differential peaks between LPC, OL NBs, and CB NBs, which highlighted additional regulators like *so* for lamina precursor cells (LPC) ^64^ and *scro* for the optic lobe progenitors ^65, 66^, while the GGG motif is detected in both CB and OL neuroblasts (Extended Data Fig. 13e). One possible candidate for this motif is the transcription factor *pros*, which is known to bind to GGG motifs and plays a major role in neuroblast/GMC identity ^59^. The addition of a temporal axis allows us to create driver lines for cell types through development. Selecting a region that is specifically open in the ONE, we created a driver line that shows a matching expression pattern in the larva and no detectable activity in the adult, thereby paralleling the decline of the neuroepithelium (Extended Data Fig. 13f).

The OL and CB neuroblasts form the roots for each of the two main branches in the developmental UMAPs (more evident in the 3D projections, Fig. 4b), indicating distinct modes of neurogenesis and maturation in each brain region. The OL shows a high amount of branching, while the CB and ventral nerve cord display a continuum, similar to our observations in the adult tSNE (Fig. 1b). Interestingly, the split between *pros*^+^ and *Imp*^+^ in central brain neurons is already present in these early stages, and, similar to scRNA-seq data, most of the ventral nerve cord neurons belong to the *Imp^+^* neurons (Fig. 4c). It has been speculated that *Imp^+^* neurons are embryonic neurons ^16^. To test this hypothesis, we used the differential regions belonging to each group and plotted their accessibility in a scATAC-seq dataset of the embryo ^58^ (Fig. 4c) Indeed, the embryonic CNS shows increased accessibility for the *Imp^+^* regions, while *pros^+^* regions only become accessible in the larval dataset. To find which transcription factors may be guiding this dichotomy, we performed motif enrichment on the differential regions. Dati and Pros motifs were enriched in the *Imp^+^* cells, most notably in peaks near *Imp* itself, suggesting a repressive role for these factors, where they may close these regions in *pros^+^* cells and suppress *Imp* expression. This layer of cell identity, related to birth order, guides most of the clustering in the central brain.

Embryonic CB neurons are rewired during early pupal stages to integrate into the adult network. This rewiring consists of a pruning step in which axons retract, followed by axonal regrowth. Both Pros and Imp have been associated with neuronal remodeling in the γ-KCs, with Imp being required for regrowth of pruned axons by promoting transport of ribonucleoprotein granules ^67, 68^, while overexpression of *pros* inhibits pruning ^69^. The process of neuronal remodeling is regulated by the ecdysone hormone and *Sox14* ^70^. We examined the regions that were specifically accessible during pruning (0h APF to 6h APF) and motif enrichment indeed reveals a spike for both EcR and Sox14 motifs (Fig. 4d). Interestingly, this spike only occurs in the *Imp^+^* neurons, suggesting that only these undergo pruning, consistent with their embryonic origin.

In the OL, six distinct branches emerge, each enriched for the motif of one major group of transcription factors (Fig. 4e). These branches show a large number of differential regions, located near genes of the immunoglobulin-like super family (median 22.5, Extended Data Table 5), suggesting a role in axonal development and synaptic partner recognition ^71, 72^. Interestingly, most of the adult OL cell types correspond to one of these branches (Fig. 4f), even though this classification does not respond to the known developmental lineages. For example, T2/T2a/T3 neurons and C2/C3 belong to the same lineage but are put in different branches ^73^. One difference we observe is in neurotransmitter usage, where cell types from branch 4 and 5 tend to be non-cholinergic. One of the eGRNs that overlaps with these branches is Ets65A, a factor previously hypothesized to be involved in non-cholinergic fate ^24^. We further note a correlation with neuropils, with most Tm neurons of the medulla and the T-neurons of the lobula plate belonging to separate branches ^73^. The cell types that have a low score for either of the branches are mostly unknown, except LLPC1 neurons that have a CB origin ^8^. Therefore, this data-driven trajectory analysis likely reflects a novel regulatory dimension with a potential role in neuronal wiring that is shared across OL neuroblast lineages.

During the cloning of eGRN regions we noticed strong differences in driver line expression between larval and adult brains. Therefore, we examined the changes of region accessibility in the eGRNs during development per cell type. Of all eGRN enhancers, 42% (3307) become more accessible in late timepoints, with 28.8% increasing after the ecdysone pulse. We also find 985 regions that undergo an enhancer switch, as their accessibility increases in one cell type while decreasing in another. One of these regions is the T1 enhancer driving *CG15117* expression shown in Fig. 3e, which is accessible in glia during development and then switches to T1 at 24h APF, with a major surge at 48h APF (Fig. 4g). Using gene expression data ^4^, we confirmed that this enhancer switch is accompanied by a gene expression switch, as *CG15117* is a glial marker during development and a T1 marker in adult. Interestingly, a small delay can be observed between enhancer accessibility and gene expression changes, similar to what was observed in other studies ^74^. Measuring the enhancer at different developmental timepoints, we indeed find GFP positive cells with the glial marker repo in development, but they disappear in the adult, coinciding with the closing of the enhancer (Fig. 4h-i).

In conclusion, by tracing back all cell types through development, we unravel a highly dynamic chromatin landscape driven by neuronal maturation and ecdysone. Large numbers of adult-specific regions are linked to reprogramming history in the CB and to neurogenesis modes in the OL. These remain accessible through development, providing a link between time points. Finally, this analysis reveals the widespread use of enhancer switching between cell types, in which specific enhancers become accessible in different cell types throughout metamorphosis.

## Discussion

We generated the first single-cell epigenome atlas of the whole fly brain throughout development, tracing neuronal and glial cell types from their birth to their mature state and providing a bridge between the regulatory code in the genome sequence and the transcriptome. Through an integrated multi-omics approach, we identified key regulators per cell type and revealed extensive gene regulatory networks. To depict this information, we introduce the concept of the enhancer-GRNs in which transcription factors are linked to high-confidence binding sites that are in turn linked to target gene expression. Given the strong insurgence of single-cell datasets, the pioneering work in single-cell ATAC-seq in mouse and human ^20, 21, 39, 75, 76^ and the fast development of single-cell multi-omics ^74, 77, 78^, eGRNs can soon be derived for many other datasets and may serve as the pinnacle for studying genomic regulatory programs.

We showed how deep learning can be integrated with omics data to accurately predict enhancer activity based on the DNA sequence. This “smarter” motif discovery approach revealed a large number of motifs that are missed by conventional algorithms and leads to a base-pair resolution prediction of binding sites. Interestingly, we noticed that some binding sites show mismatches with the canonical motif of that transcription factor, which points towards the fine-tuning of binding affinities through evolution ^79–82^.

It has long been established that neuronal identity is governed by a plethora of transcription factors, yet the study of the working mechanisms underlying these factors, namely their enhancers and target genes, has been obstructed by the complexity of the brain. There are hundreds of cell types in the fly brain, impeding the use of bulk datasets for accurately deciphering diversity. Even more, compared to the embryo ^58, 83–87^ or imaginal discs ^59, 88–94^, the number of ChIP-seq or other epigenomic datasets is very limited in the brain ^24, 95–97^. In contrast, in this study we constructed 40 eGRNs at once, covering 88 transcription factors with 7833 enhancers that are linked to 3776 target genes. Of these 88 transcription factors, 92% have lethal mutations, 72% are linked to known brain phenotypes, and 67% are linked to human diseases. Our analysis reveals that a great number of regions are regulated by multiple transcription factors, and we highlighted Ey and Mef2 as an example in Kenyon cells. The cell type specific chromatin profiles, predicted TF binding sites and interactions, and changes through development, will allow for more insight in the determination of neuronal identity.

Genetic access to neuronal cell types through the use of driver lines has revolutionized neurobiology ^11, 12, 98^. However, approaches to create driver lines have mostly been uninformed as they were based on selecting regions near genes ^99^ or randomly bashing genomic regions ^11^, leading to many unspecific driver lines. Using our scATAC-seq atlas, existing driver lines can be further dissected, and we indeed found many lines that contain multiple enhancers (median 3 and up to 9 enhancers per region in the Janelia FlyLight collection). Subcloning these individual enhancers separately, we created more specific driver lines that target distinct cell types. Furthermore, we identified 96K regions, covering 34.4% of the genome that can be queried as an initial step to create new reporter lines. Both eGRN regions and DARs can be used, the success rate increases if the region has a high correlation with gene expression. Using this system, we created new driver lines for neurons (KCs, T1, T4) and glia, and extrapolated it to development for the optic lobe neuroepithelium. Interestingly, many of the tested enhancers show increased activity during early developmental stages compared to the adult, with enhancer success rates being higher during development (Extended Data Fig. 10c). We show that this is partially caused by developmental dynamics where large numbers of enhancers open/close and switch between cell types, likely as a consequence of the high regulatory density of the *Drosophila* genome. We highlight one enhancer that switches from glial to neuronal activity, where interestingly only few transcription factors overlap. This points towards a phenotypic convergence phenomenon, where different transcription factors can regulate the same enhancer ^2^. A second reason for lower success rates in adult brains could be lower expression levels in the adult compared to developmental stages. In the embryo and imaginal discs enhancers linked directly to GFP without using the GAL4 amplification system, yield higher recall rates compared to the brain. Future enhancer-reporter assays in the brain may thus benefit from using GAL4 as reporter system. Some enhancers may furthermore depend on a specific proximal promoter. Nevertheless, we believe that success rates of >40% are high enough to create a driver line for any cell type. Our atlas also opens up the possibility of creating temporo-controlled driver lines using the split-GAL4 system with enhancers corresponding to different maturation modules.

Although for the central brain, we have lower cell type resolution in scATAC-seq compared to scRNA-seq, we detected large overarching programs between *pros*^+^ and *Imp*^+^ cells. We are able to trace this duality back to the embryo, where only *Imp*^+^ cells exist and show that these neurons are reprogrammed during metamorphosis, confirming previous hypotheses ^16^. Future studies including single-cell multi-ome assays will likely resolve additional cell types in the central brain. Finally, we detect six different modes of neurogenesis in the optic lobe, each dominated by a specific TF family that we can link to neuronal wiring and neurotransmitter expression.

We believe that the regulatory atlas of the brain, covering cell types, transcription factors, and enhancer regions, together with their joint representation as eGRNs, will be of great value to the community. Therefore, we have made all the data generated in this project publicly available at http://flybrain.aertslab.org. The website allows users to explore regulatory networks of key transcription factors with target regions and genes, and links out to SCOPE (http://scope.aertslab.org/#/Fly_Brain/) and the UCSC Genome Browser (http://genome.ucsc.edu/cgi-bin/hgTracks?db=dm6&hubUrl=http://ucsctracks.aertslab.org/papers/FlyBrain/hub.txt).

## Extended data figures

**Extended data Fig. 1:**
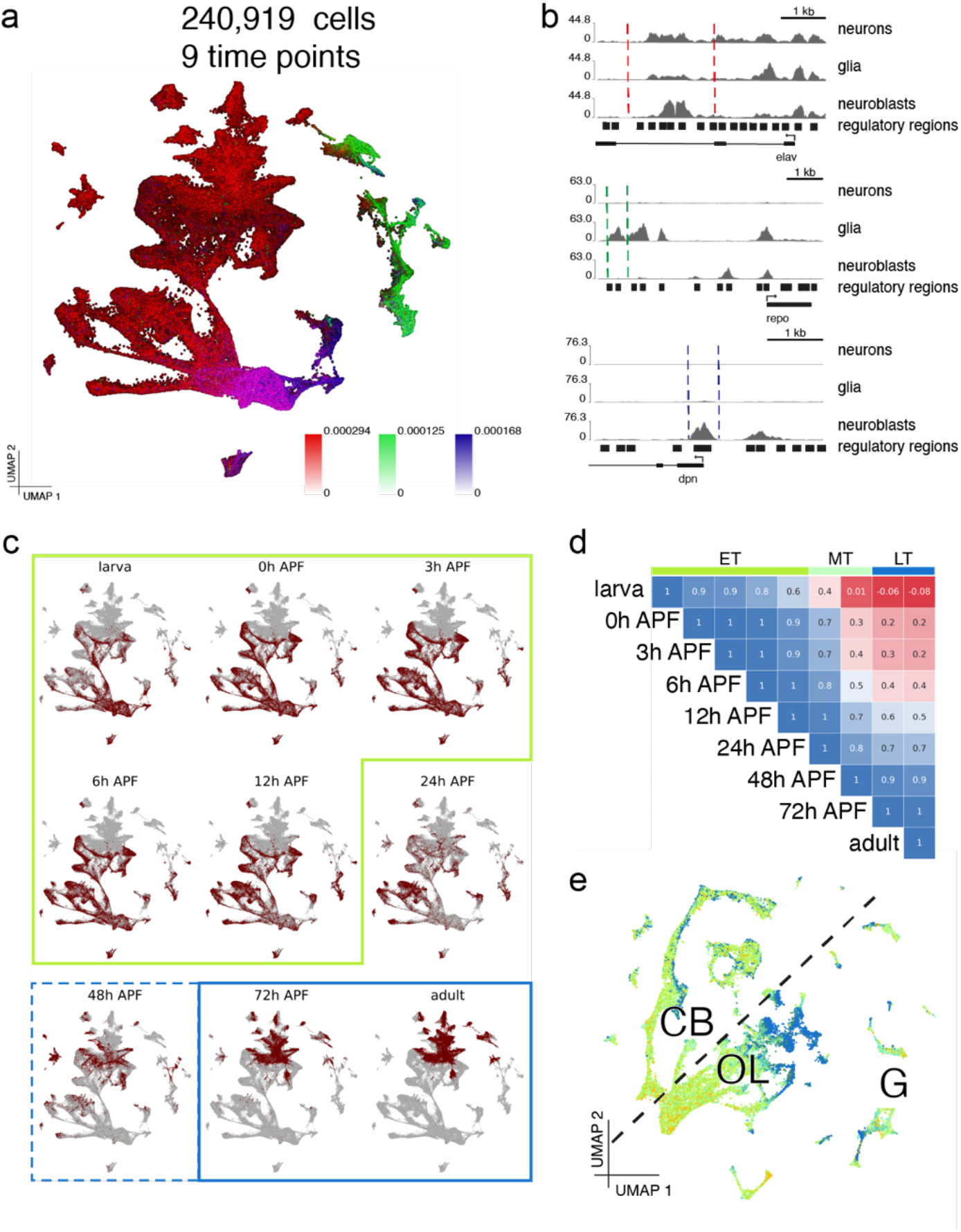
Global analysis of all timepoints reveals major dynamics and optic lobe-central brain split. **a.** UMAP of global cisTopic (150k cells shown), colored by region accessibilities near elav (red), repo (green) and dpn (blue), matching neurons, glia and neuroblasts respectively **b.** Overview of regions shown in **a.** for representative cell types (Kenyon cells for neurons, Astrocytes for glia, optic lobe neuroblasts for neuroblasts) **c.** Distribution of cells per timepoint in the global UMAP; The border groups timepoints jointly analyzed in the upcoming sections (green: early timepoints, blue: late timepoints) **d.** Spearman correlation of top 1000 variable regions across timepoints **e.** UMAP after timepoint correction with Harmony (colored by timepoint: early timepoints in green, late in blue).

**Extended data Fig. 2:**
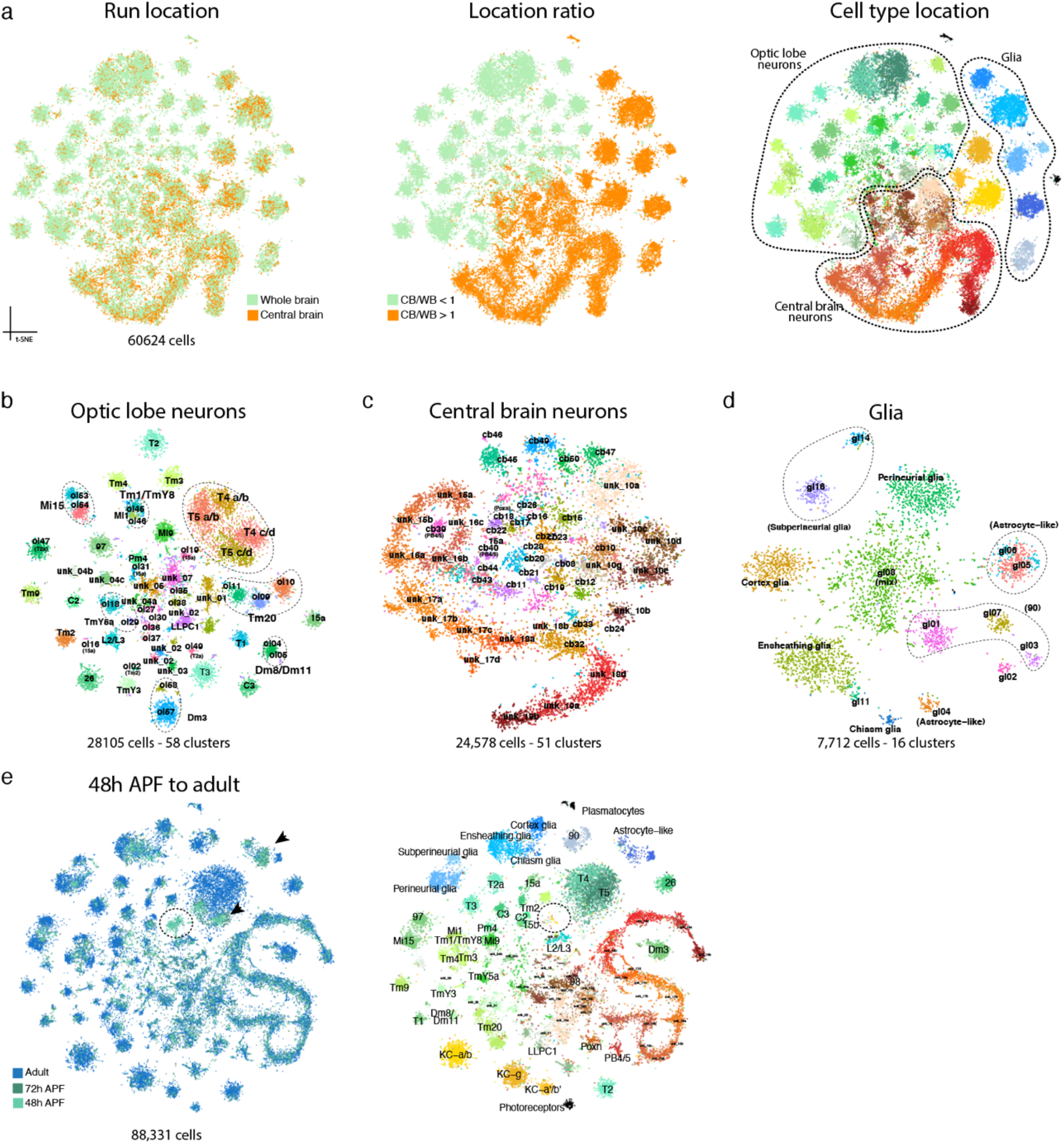
Sub-clustering of main categories in the adult. **a.** t-SNE of the 60k cells in the adult cell types analysis (includes adult and 72h pupa). The different colouring schemes illustrate how the enchment of central brain only runs leads to the annotation of the cell clusters according to their location (central brain and optic lobe). Sub-clustering of the cells was performed splitting the neurons based on their location and glia, note that Kenyon cells, plasmatocytes and photoreceptors were not included in these major groups. **b.** Sub clustering of optic lobe neurons leads to 58 sub clusters, including a further split of T4/T5 neurons. **c.** Sub clustering of central brain neurons reveals 51 subtypes. Notice how the S-shape of Pros+ cells and the split from Imp+ cells are retained. **d.** Sub clustering of glia reveals 16 subtypes. **e.** Clustering including 88k cells from 48h APF to adult, provides three extra clusters including younger cells, but does not increase the resolution of the adult cell types.

**Extended data Fig. 3:**
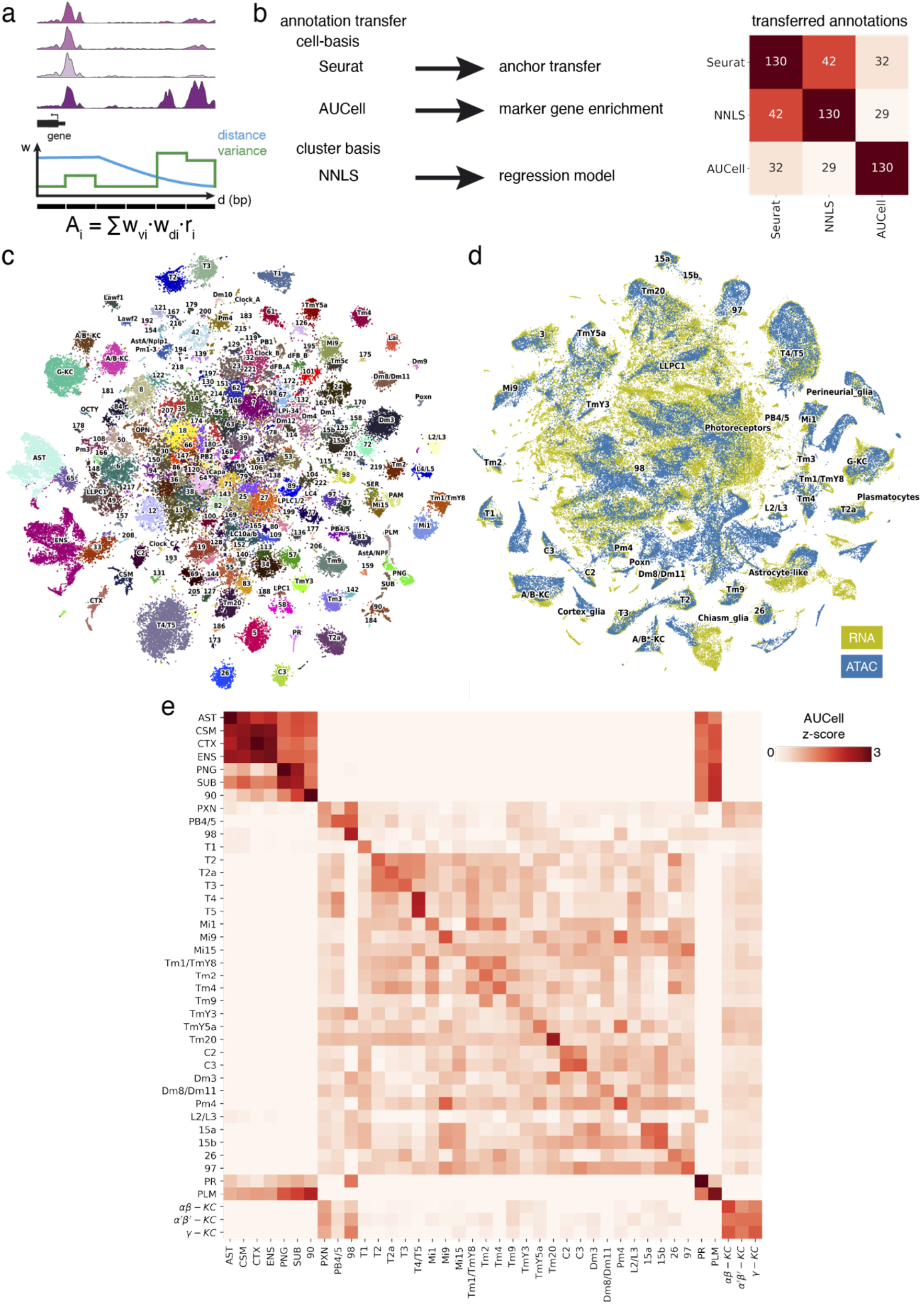
Integration of scRNA-seq and snATAC-seq. Calculation of gene-accessibility scores using a weighted sum of regions in the gene body and up to 5kb upstream of the TSS. Weights decrease exponentially with distance from the TSS (constant in the gene body), and increase with higher gini coefficients **b.** Overview of annotation methods used. Main cell types are detected with each method, while low confidence matches are method specific **c.** Annotated tSNE of the transcriptomes of 118k adult cells **d.** Integrated tSNE of scRNA-seq and snATAC-seq **e.** Gene set enrichment with of marker genes using AUCell on gene-accessibility matrix, revealing matches per cell type and per major cell type group (glia, optic lobe neurons, central brain neurons, Kenyon cells, photoreceptors and plasmatocytes).

**Extended data Fig. 4:**
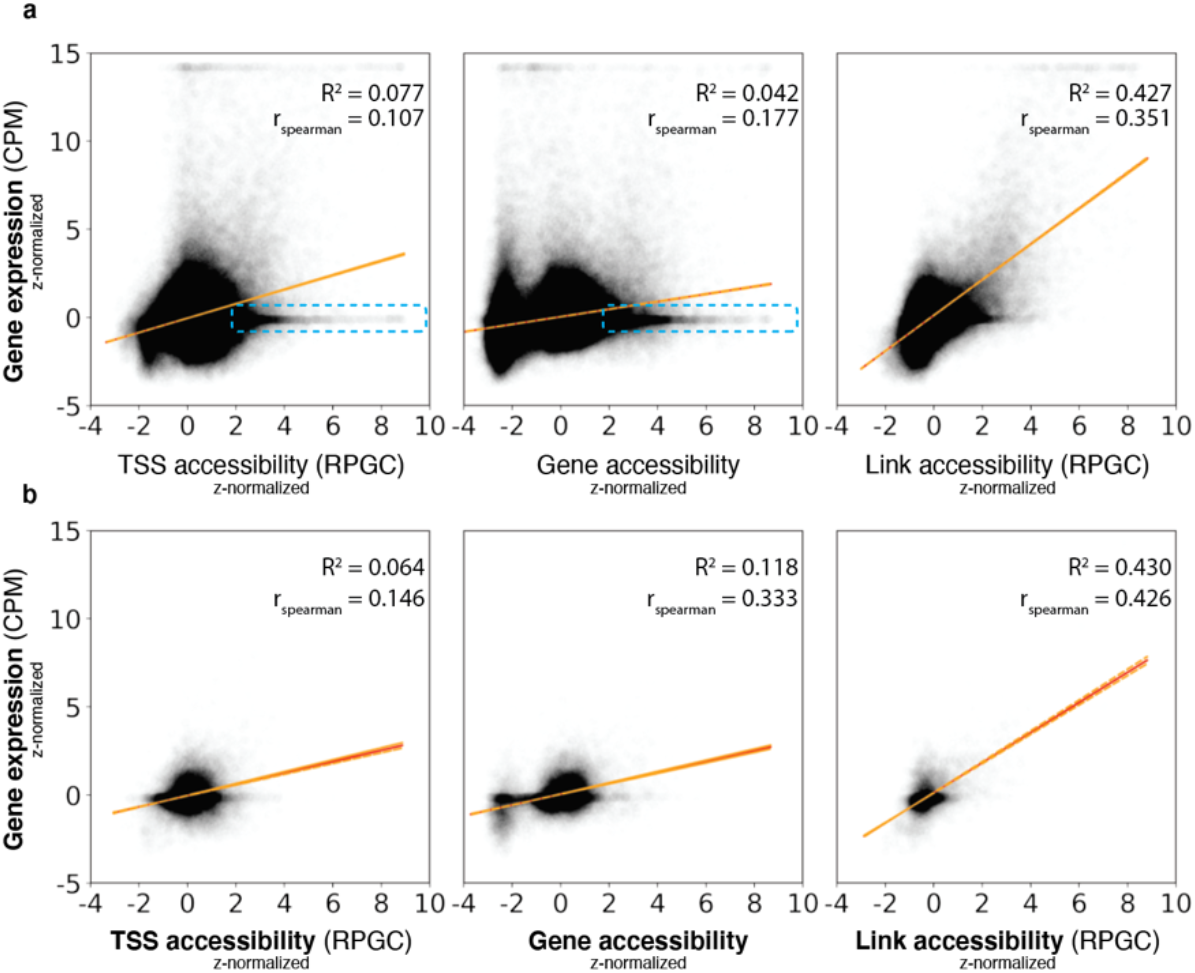
Gene expression correlates with region accessibility. Correlation of gene expression (**a.**) and TF gene expression (**b.**) with aggregated accessibility profiles at TSS, averaged gene-accessibility, and averaged accessibility of regions with positive links. Red line shows linear fit, with orange boundaries as the 95% confidence interval. Note the regions near the TSS that have high accessibility but do not lead to gene expression (highlighted in blue) and the increase in performance for the gene-accessibility score in the TF expression, while overall highest correlation is reached with links.

**Extended data Fig. 5:**
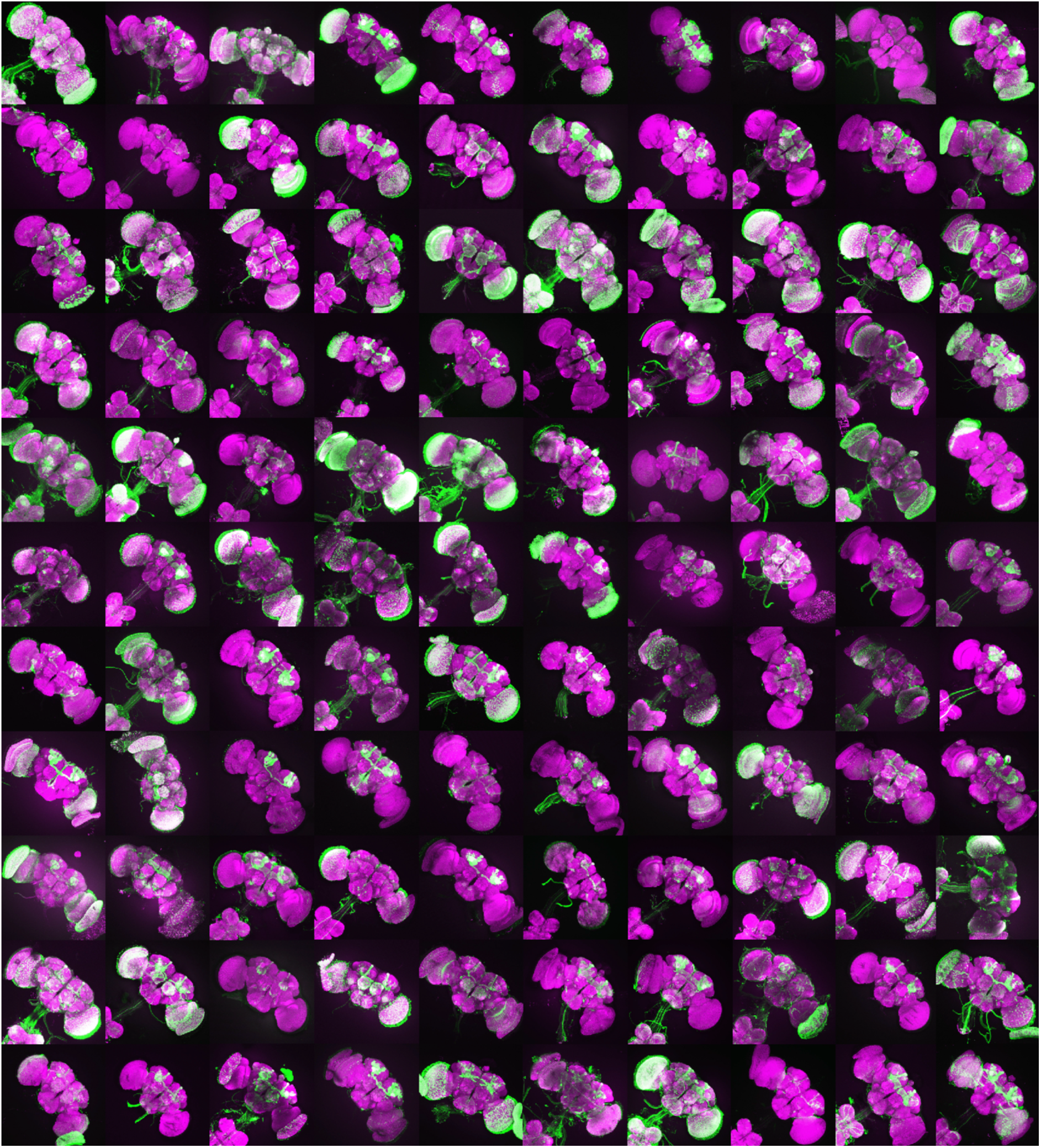
Cell type specific regions can serve as functional enhancers. Kenyon cell DARs overlap with 588 Janelia regions active in Kenyon cells, of which a subset of 110 is shown here. Images courtesy of the Janelia FlyLight Project.

**Extended data Fig. 6:**
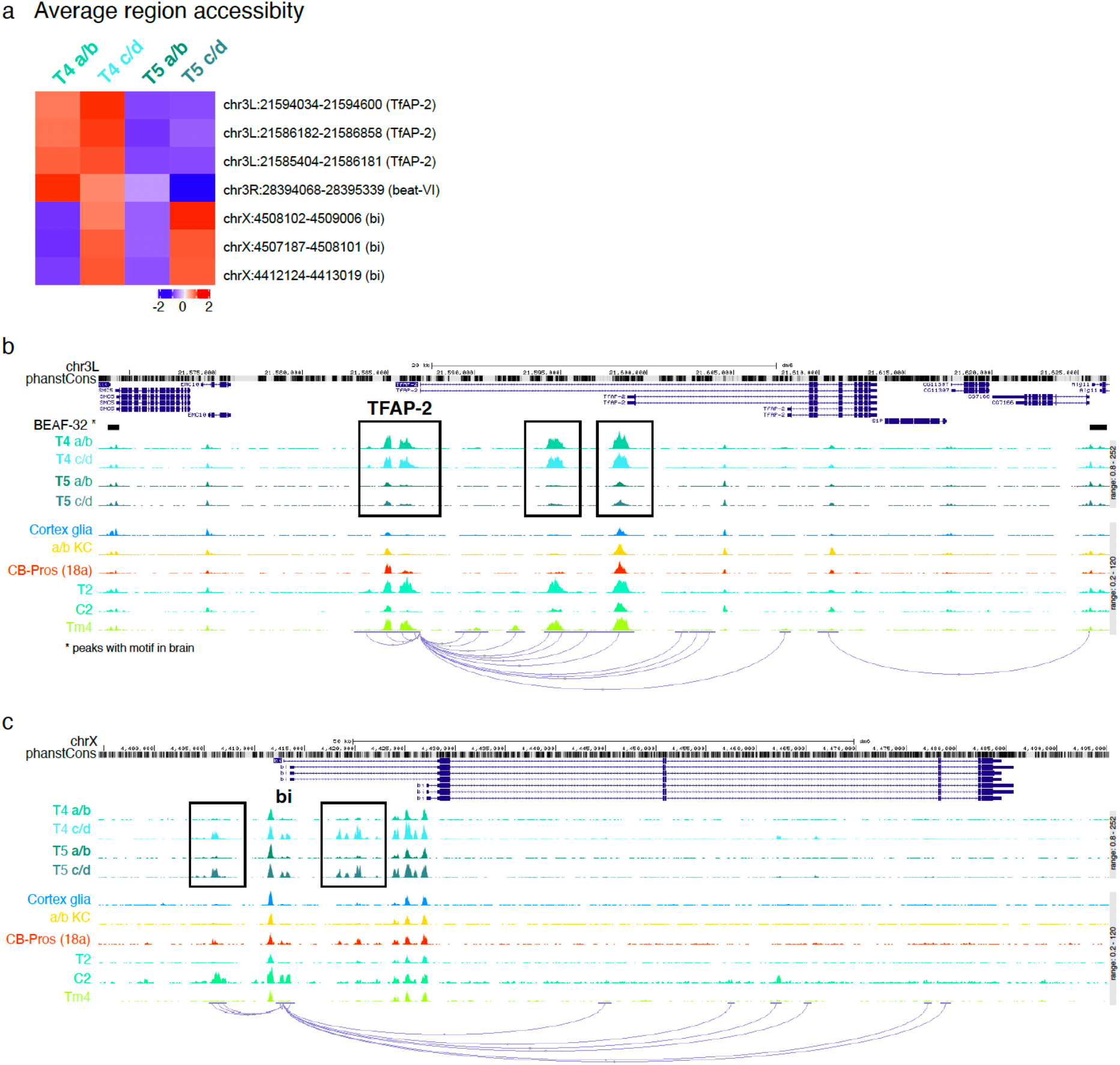
Identification of T4/T5 subtypes. Subclustering of T4 and T5 neurons identifies the a/b and c/d subtypes, with differential regions near marker genes *TfAP-2* and *bi*. **b.** Locus of *TfAP-2*, showing differential peaks between T4 and T5 neurons **c.** Locus of *bi* shows specific peaks for c/d subtype.

**Extended data Fig. 7:**
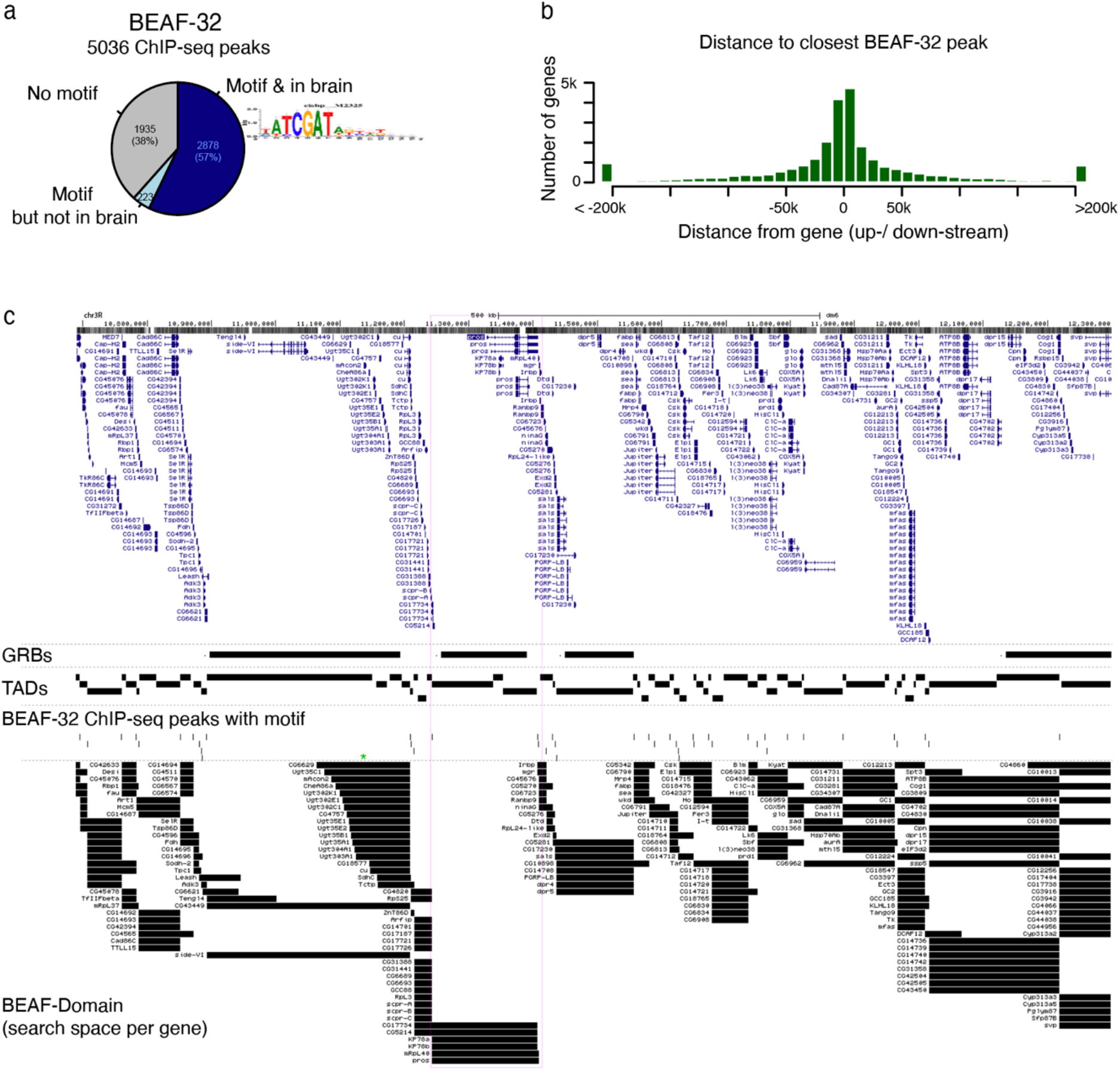
BEAF-32 ChIP-seq peaks can be used to delimit the search space for regulatory regions around each gene. **a.** Proportion of BEAF-32 ChIP-seq peaks that have a high-scoring BEAF-32 motif, and are accessible in the fly brain. Despite being performed on whole embryo (0-14 hours, mixed sex), most of the peaks with motif are ubiquitously accessible across the cell types in the brain. **b.** Distance to the closest BEAF-32 peak with motif upstream and downstream of each gene. Most of the genes (86%) are within 50kbp of a BEAF-32 peak (46% are between two peaks within 50kbp, and 88% within 200kbp). For those genes further than 50kbp, expanding the search space from 50kbp to 200kb adds a median of 2 extra links. **c.** View of genomic regulatory blocks (GRBs), topological associated domains (TADs) and BEAF-32 ChIP-seq peaks near *pros* (highlighted by the pink rectangle). For the genes within the defined genomic regulatory blocks, 95% of these associations are contained within the same block, and 25% within the same TADs. Since the current GRBs dataset does not provide enough genome coverage (it only includes 1523 genes, 15% of the 9821 genes with links), and the TADs are very fragmented (only 67% of genes have their biggest transcript within one TAD), we opted to use the BEAF-32 peaks to define the search space per gene. The lowest track shows the search space used for each gene (i.e. the region between the first two BEAF-32 peak within 200kbp of the transcript, skipping the 500bp around the TSS. In case there are no peaks within 200kbp, 50kbp is kept as search space).

**Extended data Fig. 8:**
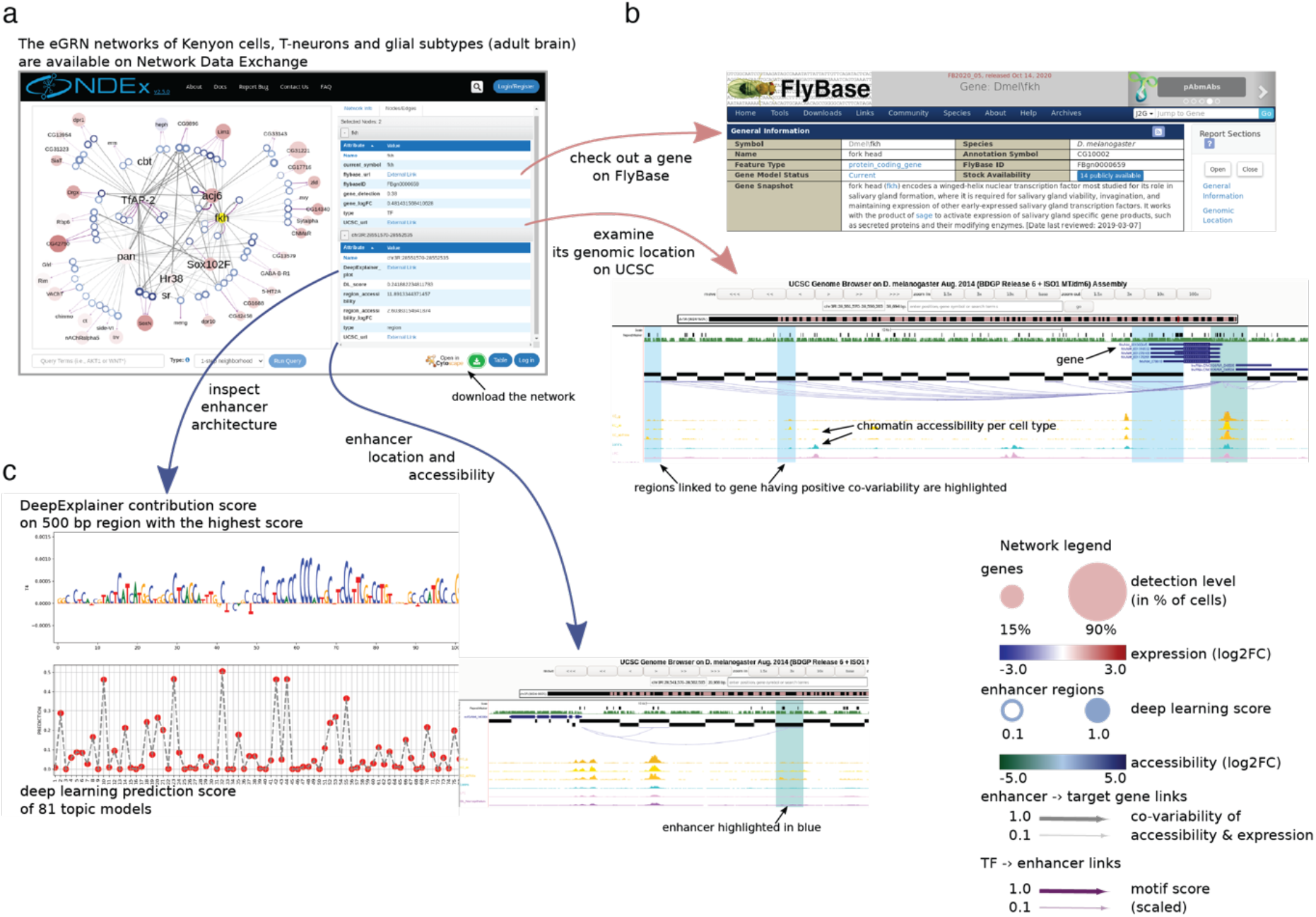
eGRN overview. **a.** eGRNs were determined for different subytpes of Kenyon cells, T-neuron subclasses and glia and are available for exploration on NDEx through https://flybrain.aertslab.org/. **b.** Link outs from the gene to FlyBase and UCSC allow to explore gene function and chromatin profiles with all nearby predicted enhancers colored. **c.** Link outs from regions allow to inspect the region with DeepFlyBrain, to visualize nucleotide importances, while also linking to UCSC to view the genomic context with the selected region highlighted.

**Extended data Fig. 9:**
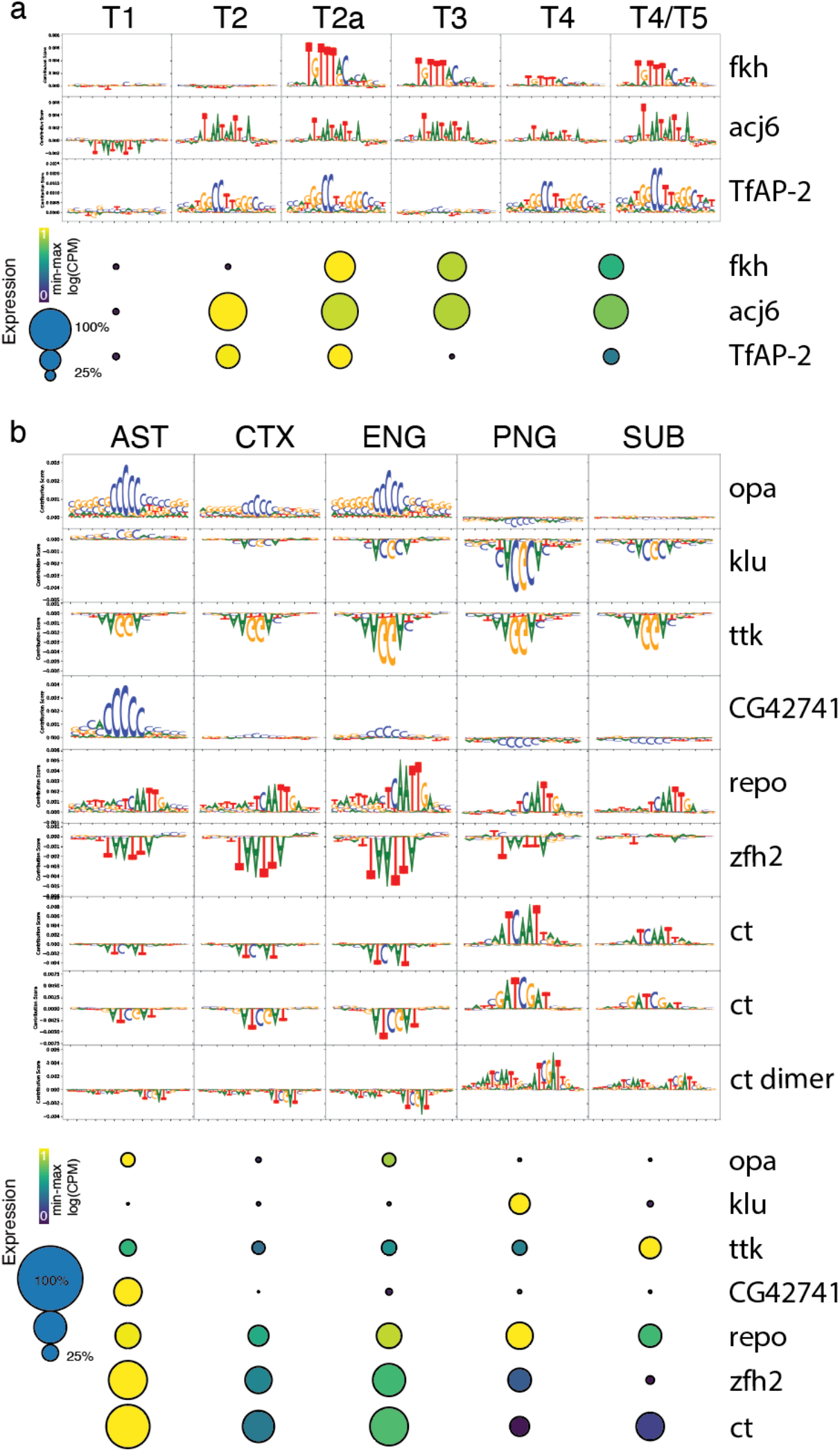
Deep learning predicts de novo key transcriptional activators and repressors. **a.** Motifs used in the convolutional filters of the model to classify T-neuron regions predict Fkh, Acj6 and TfAP-2 as activators in cell types matching their expression profiles. **b.** Motifs identified by the model to classify glial regions reveals activators (Opa, CG42741, Repo and Ct) and repressors (Klu, Ttk, Zfh2), matching expression profiles.

**Extended data Fig. 10:**
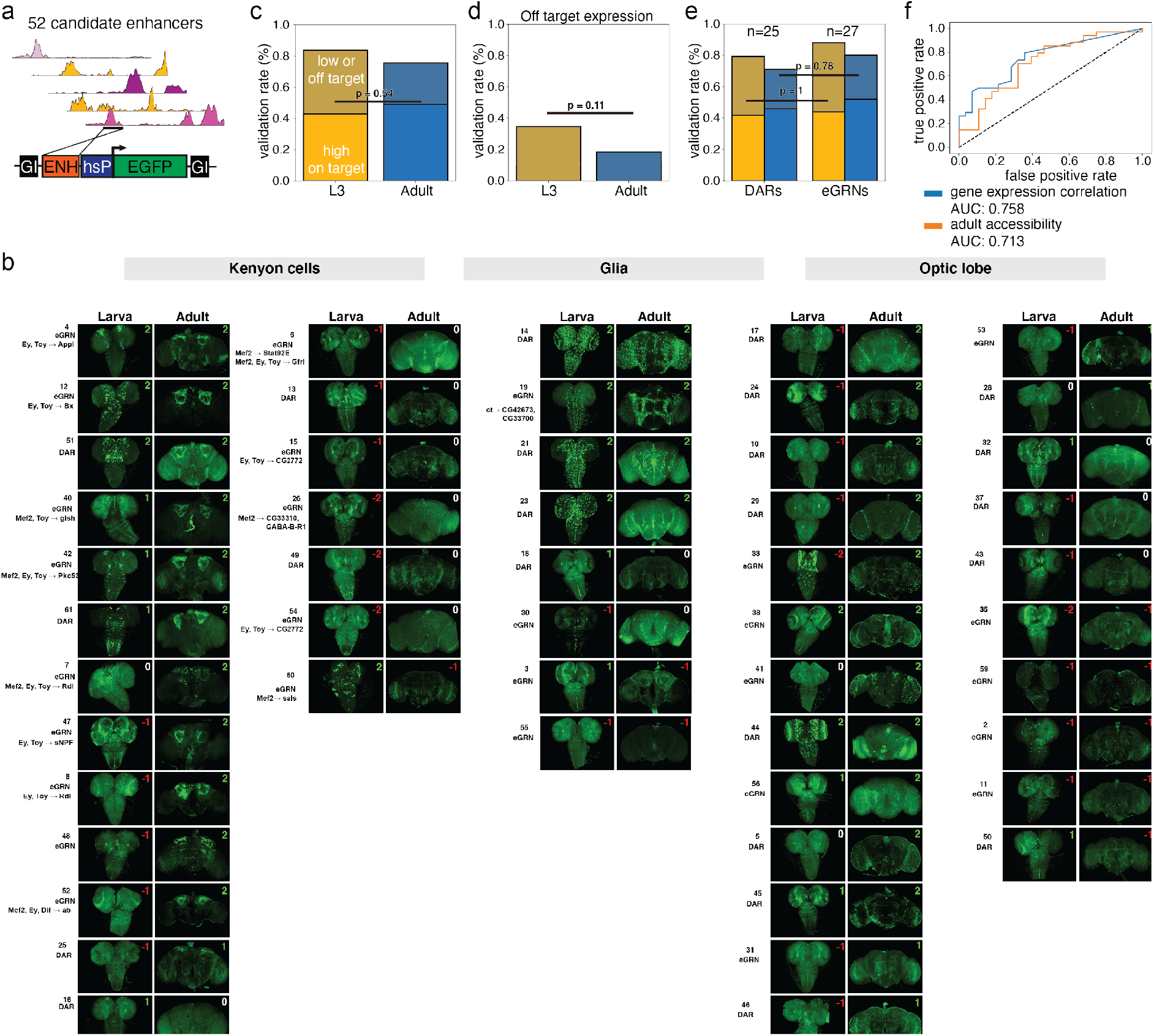
Enhancers selected by DARs or eGRNs generate novel driver lines. **a.** 52 enhancers were selected and cloned into a construct, flanked by gypsy insulators (GI) driving GFP expression from the Hsp70 promotor. Selected peaks have a median size of 579bp. **b.** Overview of GFP expression in different cloned enhancers. **c.** Bar plots showing validation rate for GFP expression with a distinction for high on target expression vs low or off target expression. Success rates are higher in the larval brain compared to the adult (two-sided Fischer’s exact, n=52). **d.** Off target expression reaches 40 and 20% in larval and adult brains respectively (two-sided Fischer’s exact, n=52). **e.** Both DARs and eGRN regions can be used to create driver lines (two-sided Fischer’s exact, n shown). **f.** Precision-recall curve shows accessibility and correlation of region with nearby gene expression are key features to predict GFP activity.

**Extended data Fig. 11:**
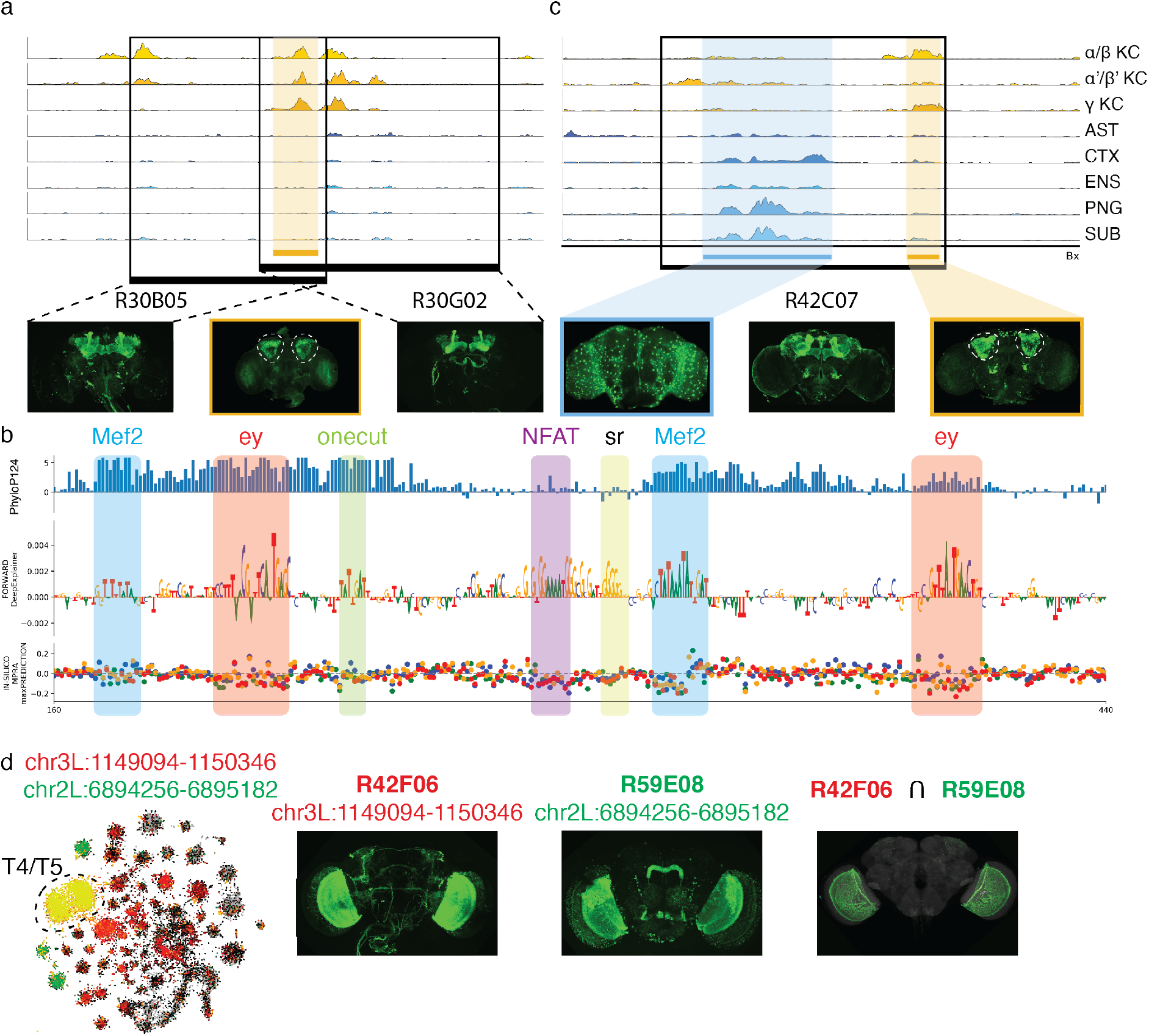
Breakdown of existing driver lines into functional components. **a.** Peak in the overlap of two existing KC driver lines recapitulates KC expression. **b.** DeepExplainer view of the functional element from (a) showing ey and Mef2 binding sites. **c.** Existing non-specific driver lines can be broken in separate more specific drivers for Kenyon cells and glia using cell-type specific ATAC-peak signals. **d.** In-silico overlap of ATAC-peak signals resembles that of in-vitro split-GAL4 lines for T4/T5 neurons. Images courtesy of the Janelia FlyLight Project.

**Extended data Fig. 12:**
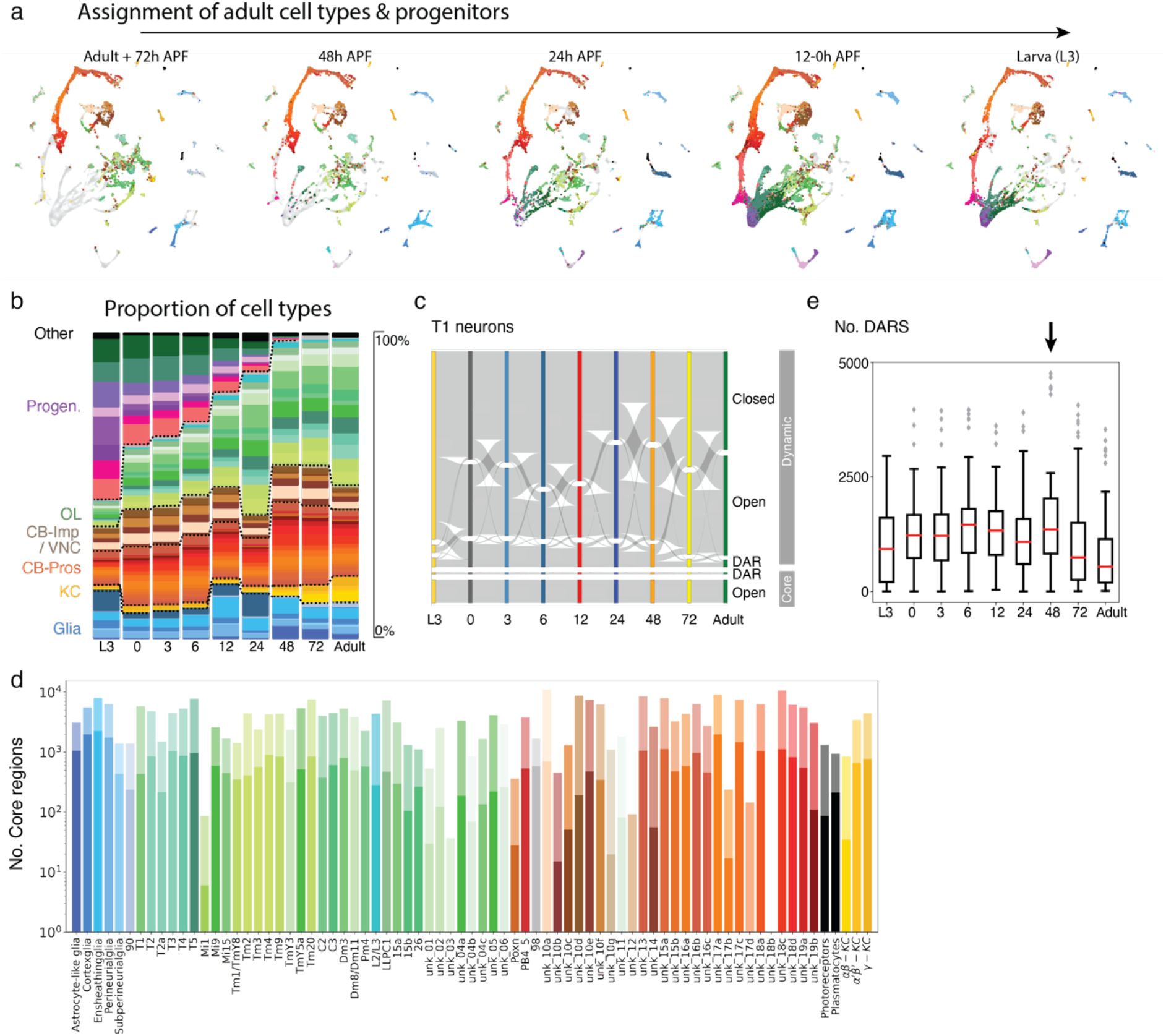
Tracking cell types across development reveals presence of core-regions. **a.** An SVM classifier was used to propagate the adult cell type labels to earlier stages in development. The classifier also included the progenitor cell types –from the developmental analysis– (purple and dark green colors), to provide further confidence. **b.** Proportion of cells with each label at each developmental timepoint. **c.** Chromatin landscapes for T1 neurons, shows a highly dynamic opening and closing of peaks during development. A core set remains accessible at all times, of which a subset is specific to T1 neurons. **d.** Bar plots showing the number of core-regions identified per cell type. Dark colors show specific core regions (core-DARs). **e.** Number of DARs calculated per cell type (down sampled to 75 cells) for every timepoint shows a decline over time. The arrow notes a small increase at 48h APF during synaptogenesis. Red line highlights the median, 25th and 75th percentiles are shown as the box edges, and data points within 1.5 times the interquartile range from the edge (whiskers) and outliers are shown as data points, n=74,77,78,77,78,77,78,75,76 cell types.

**Extended data Fig. 13:**
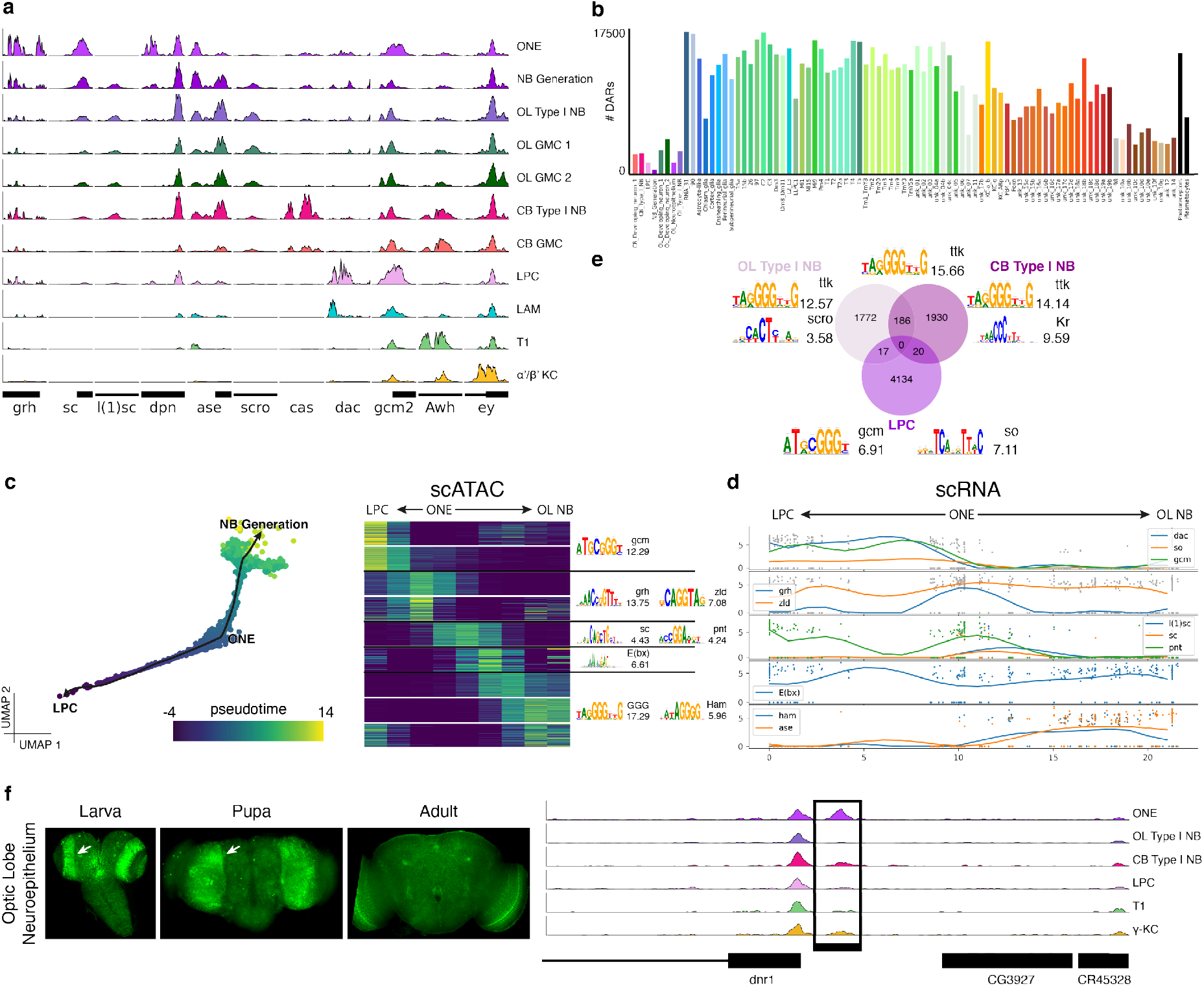
Neural progenitors have unique chromatin profiles. **a.** Progenitor cell types show specific marker accessibility, while neurons already show accessibility in adult specific regions **b.** Number of DARs per cell type in the early development dataset, revealing a lower number for progenitors (purple shades) **c-d.** Trajectory from optic lobe neuroepithelium (ONE) to lamina progenitor cells (LPC) and optic lobe neuroblasts (OL NB) using scATAC-seq (**c**) and scRNA-seq (**d**). Heatmap shows dynamic chromatin accessibility modules with enriched motifs (NES score shown) and line plot shows expression profiles for predicted master regulators. **e.** Specific comparison of different progenitor cell types detects thousands of differential regions, with enrichment of motifs of key transcription factors **f.** GFP reporter showing activity of ONE specific region in early pupal timepoints, followed by a decrease in signal.

## Methods

### Data reporting

No statistical methods were used to predetermine sample size. The experiments were not randomized and the investigators were not blinded to allocation during experiments and outcome assessment.

### Genetics

Flies were raised on a yeast-based medium at 25°C on a 12h/12h day/night light cycle. All *Drosophila* lines used in the single-cell ATAC-seq experiments are derived from the DGRP collection. One hybrid was created by crossing different DGRP lines, generating genetic diversity.

| *Drosophila* line | Source | Identifier | Genetics |
| --- | --- | --- | --- |
| *D. melanogaster*: DGRP-551 | Bloomington Drosophila Stock Center | BDSC: 55026;<br>FlyBase: FBsn0000297 | Wild type |
| *D. melanogaster*: DGRP-409 | Bloomington Drosophila Stock Center | BDSC: 28278;<br>FlyBase: FBsn0000111 | Wild type |
| *D. melanogaster*: DGRP-502 | Bloomington Drosophila Stock Center | BDSC: 28204;<br>FlyBase: FBsn0000124 | Wild type |
| *D. melanogaster*: DGRP-639 | Bloomington Drosophila Stock Center | BDSC: 25199;<br>FlyBase: FBsn0000141 | Wild type |
| *D. melanogaster*: Hybrid (DGRP-551, DGRP-360, DGRP-907, DGRP-913) | This lab |  | Wild type |
| *D. melanogaster*: MB371B | Bloomington Drosophila Stock Center | BDSC: 68383 | w[1118];<br>Py[+t7.7]<br>w[+mC]=R13F02-p65.ADattP40;<br>Py[+t7.7]<br>w[+mC]=R85D07-GAL4.DBDattP2 |
| *D. melanogaster*: MB418B | Bloomington Drosophila Stock Center | BDSC: 68322 | w[1118]; Py[+t7.7]<br>w[+mC]=R26E07-p65.ADattP40/CyO<br>Py[+t7.7]<br>w[+mC]=R30F02-GAL4.DBDattP2 |
| *D. melanogaster*: MB419B | Bloomington Drosophila Stock Center | BDSC: 68323 | w[1118];<br>Py[+t7.7]<br>w[+mC]=R26E07-p65.ADattP40/CyO:<br>Py[+t7.7]<br>w[+mC]=R39A11-GAL4.DBDattP2 |
| *D. melanogaster*: UAS-GFP.nls14 | Bloomington Drosophila Stock Center | BDSC: 4775 | w[1118];<br>Pw[+mC]=UAS-GFP.nls14 |
| *D. melanogaster*: UAS-GFP.nls8 | Bloomington Drosophila Stock Center | BDSC: 4776 | w[1118];<br>Pw[+mC]=UAS-GFP.nls8 |

### Nuclei isolation

*D. melanogaster* adult brains were dissected and transferred to a tube containing 100 µl ice cold DPBS solution. After centrifugation at 800 g for 5 min, the supernatant was replaced by 500 µl nuclei lysis buffer composed of 10 mM Tris-HCl pH 7.4, 10 mM NaCl, 3 mM MgCl2, 0.1% Tween-20, 0.1% Nonidet P40, 0.01% Digitonin, and 1% BSA, in Nuclease-free water. The following procedure was followed to extract the nuclei from the brain tissue: incubation in nuclei lysis buffer on ice for 5 min, transfer to a dounce tissue grinder tube (Merck), 25 strokes with pestle A, incubation on ice for 10 min, 25 strokes with pestle B. The lysis was stopped by adding 1 ml of wash buffer composed of 10 mM Tris-HCl pH 7.4, 10 mM NaCl, 3 mM MgCl2, 0.1% Tween-20 and 1% BSA, in Nuclease-free water. Nuclei were pelleted by centrifugation at 800 g for 5 min at 4°C and resuspended in 1x Nuclei Buffer (10x Genomics). Nuclei suspensions were passed through a 40 µM Flowmi filter (VWR Bel-Art SP Scienceware). Nuclei concentration was assessed by the LUNA-FL Dual Fluorescence Cell Counter.

### 10x Genomics

Single-cell libraries were generated using the GemCode Single-Cell Instrument and Single Cell ATAC Library & Gel Bead Kit v1 and ChIP Kit (10x Genomics, US). Briefly, fly brain single nuclei were suspended in 1x nuclei buffer. The single nuclei were incubated for 60 min at 37°C with a transposase that fragments the DNA in open regions of the chromatin and adds adapter sequences to the ends of the DNA fragments. After generation of nanoliter-scale Gel bead-in-Emulsions (GEMs), GEMS were incubated in a C1000 Touch Thermal Cycler (Bio Rad) programmed at 72°C for 5 min, at 98°C for 30 s, 12 cycles of (98°C for 10 s, 59°C for 30 s, 72°C for 1 min), and held at 15°C. After incubation, single-cell droplets were broken and the single-strand DNA was isolated and cleaned with Cleanup Mix containing Silane Dynabeads. Illumina P7 sequence and a sample index were added to the single-strand DNA during library construction via PCR: at 98°C for 45 s, 11-13 cycles of (98°C for 20 s, 67°C for 30 s, 72°C for 20 s), 72°C for 1 min, and hold at 4°C. The sequencing-ready library was cleaned up with SPRIselect beads.

### Sequencing

Before sequencing, the fragment size of every library was analyzed on a Bioanalyzer high-sensitivity chip. All 10x scATAC libraries were sequenced on NextSeq500 and NovaSeq6000 instruments (Illumina) with the following sequencing parameters: 50 bp read 1 – 8 bp index 1 (i7) – 16 bp index 2 (i5) – 49 bp read 2.

### Omni-ATAC-seq of FAC-sorted samples

100 GFP-expressing (MB371B, MB418B, MB419B crossed with UAS-nls.GFP) and 15 GFP negative (wild type) fly brains were dissected in PBS on ice. The brains were then centrifuged at 800 g for 5 min, after which the supernatant was replaced by 50 μL of dispase (3 mg/mL, Sigma-Aldrich_D4818-2mg), 75 μl collagenase I (100 mg/mL, Invitrogen_17100-017), and 125 μL trypsin-EDTA (0.05%, Invitrogen_25300054). Brains were dissociated at 25°C in a Thermoshaker (Grant Bio PCMT) for 15 min at 25°C at 1,000 rpm and the solution was mixed by pipette every 5 min. After, cell suspensions were passed through a 10 μM pluriStrainer (ImTec Diagnostics_435001050) and viability was assessed by the LUNA-FL Dual Fluorescence Cell Counter. Next, 4 aliquots were made containing GFP-brains cells with/without PI (10%) and GFP-positive brains with/without PI (10%). FACS was performed on the FACS Aria III (BD Biosciences, US). The GFP-negative brains were used to set the gates on the machine for cell size and viability (PI), the GFP-positive brains for the GFP fluorescence, after which the GFP positive cells with PI were sorted (see Supplementary Data 1). Between 2.6 and 11k GFP positive cells were sorted and 50k cells per negative control. After sorting, regular omni-ATAC-seq was performed as described by Corces et al. ^38^.

### Immunohistochemistry

For immunofluorescence, brains were dissected and transferred to a tube containing 100 μl ice cold DPBS solution. After centrifugation at 800 g for 5 min, the supernatant was replaced by 4% Formaldehyde in PBT 0.3% (DBPS + 0.3% Triton X-100 (Sigma)) and incubated at room temperature with rotation for 15 min. Brains were washed with PBT 0.3% three times, rotating for 10 min at room temperature each time and then blocked in Pax-DG (10 g BSA (Sigma), 3g Deoxycholate Acid (Sigma), 3ml Triton X-100 (Sigma), 50ml Normal Goat Serum (MP Biomedicals), 100ml 10X PBS, 850ml H2O) for 2 hours at room temperature with rotation. Primary antibody mixes were created in Pax-DG (dilutions detailed in the Key Resources Table) and brains were incubated in these mixes overnight at 4°C with rotation. The next day, brains were washed with PBT 0.3% three times, rotating for 10 min at room temperature each time and then stained with secondary antibody mixes in Pax-DG (dilutions detailed in the Key Resources Table) for 2 hours at room temperature with rotation. Brains were washed with PBT 0.3% three times, rotating for 10 min at room temperature each time and mounted in Mowiol mounting medium (Sigma). Imaging was performed using Nikon C2 and Nikon A1 confocal microscopes.

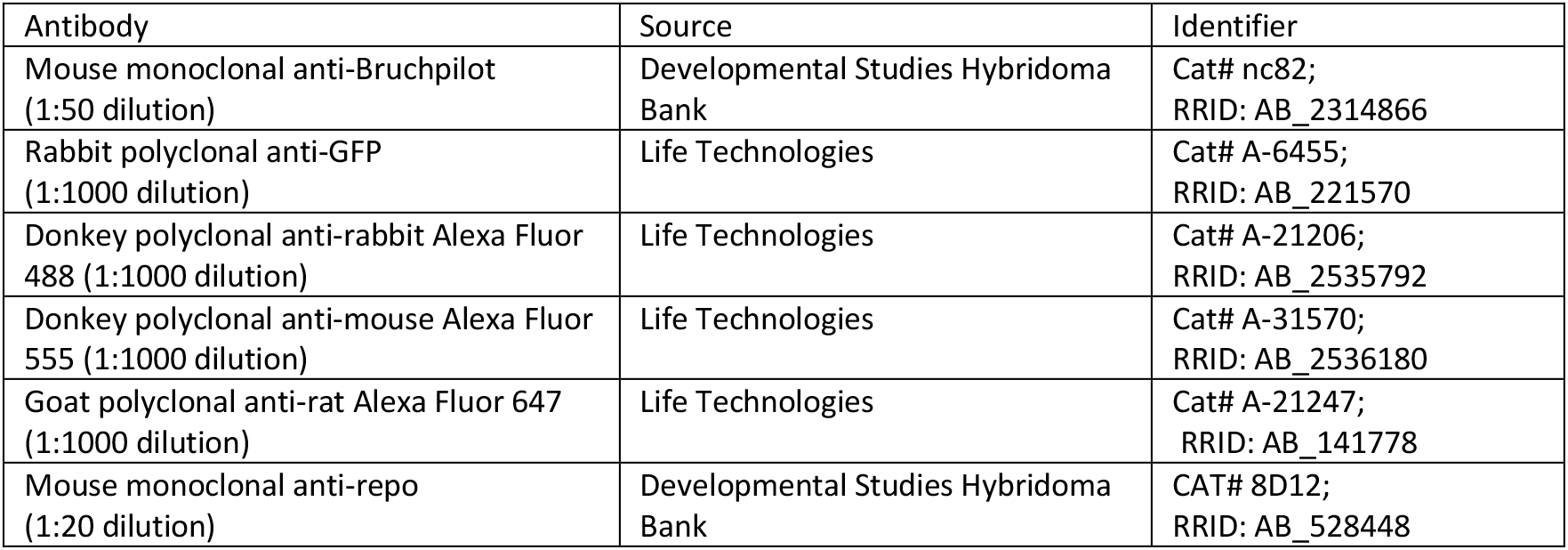

| Antibody | Source | Identifier |
| --- | --- | --- |
| Mouse monoclonal anti-Bruchpilot (1:50 dilution) | Developmental Studies Hybridoma Bank | Cat# nc82;<br>RRID: AB_2314866 |
| Rabbit polyclonal anti-GFP (1:1000 dilution) | Life Technologies | Cat# A-6455;<br>RRID: AB_221570 |
| Donkey polyclonal anti-rabbit Alexa Fluor 488 (1:1000 dilution) | Life Technologies | Cat# A-21206;<br>RRID: AB_2535792 |
| Donkey polyclonal anti-mouse Alexa Fluor 555 (1:1000 dilution) | Life Technologies | Cat# A-31570;<br>RRID: AB_2536180 |
| Goat polyclonal anti-rat Alexa Fluor 647 (1:1000 dilution) | Life Technologies | Cat# A-21247;<br>RRID: AB_141778 |
| Mouse monoclonal anti-repo (1:20 dilution) | Developmental Studies Hybridoma Bank | CAT# 8D12;<br>RRID: AB_528448 |

### 10x data processing

The 10x fly brain samples were each processed (alignment, barcode assignment and UMI counting) with CellRangerATAC 1.2.0 count pipeline. The Cell Ranger reference index was built upon the 3rd 2017 FlyBase release (D. melanogaster r6.16) ^100^. Sequencing saturations were calculated based on Michaelis-Menten kinetics and early pupal timepoints were additionally sequenced and CellRanger aggr was used to aggregate sequencing results.

### Demuxlet

We used Demuxlet ^101^ to demultiplex the different genotypes that were used in the DGRP-mixed samples, allowing us to remove doublets of two different genetic backgrounds. The vcf file of the DGRP project (available at http://dgrp2.gnets.ncsu.edu/) was lifted over to dm6 genome and SNPs for DGRP-409 and DGRP-502 were extracted. For DGRP-639 and the DGRP-551 based hybrid we performed bulk ATAC-seq to generate updated SNP profiles. After combining all SNPs, we only retained SNPs that were unique for one line. This vcf file was then used in Demuxlet with default parameters leading to the identification and removal of 43,489 doublets (details in Extended Data Table 1).

### scATAC topic modelling

After removing doublets, we performed some extra QC filters to select the 240,919 cells that will be used in upcoming analyses (Signac’s nucleosome_signal <= 10, global blacklist_ratio < 0.05 and non-outlier blacklist ratio within its own run, number of fragments between 100 and 50k).

To run cisTopic ^34^, we created the cell-counts matrix using 129,109 pre-defined regulatory regions (ctx) based on conservation ^102^ (i.e. counted fragments within these regions). Given the large size of our dataset, we implemented WarpLDA ^103^ within the cisTopic package as a faster and more efficient alternative to Collapsed Gibbs Sampling (CGS). WarpLDA uses delayed update approach, meaning that topic-region and cell-topic distributions are updated after a number of assignments rather than after each assignment, reducing the number of calculations and memory access. This new faster algorithm is now available in cisTopic version 3 (https://github.com/aertslab/cisTopic).

We performed topic modeling on the whole matrix (with between 2 and 500 topics, with 500 iterations, and finally selecting the model with 500 topics). This analysis was used to obtain an overview of the whole dataset, and to perform the analyses across development. However, we noticed that we obtained slightly better region accessibility predictions, and higher clustering resolution, when analyzing subsets of the dataset (e.g. the T4/T5 split is not detected in this global analysis, the TfAP-2 enhancers are not predicted as differential). Therefore, we used independent cisTopic runs to perform the analysis of the adult cell types (including adult and 72h APF, using 200 topics), and developmental stages (larva and 0-12h APF, 200 topics). This split of stages was chosen based on their similarity (e.g. Extended Data Fig. 1c,d).

Adult cell clusters were defined based on a two-level analysis: (1) First, on the “Adult + 72h APF” cisTopic run, we clustered the adult cells –with more than 900 fragments in peaks (FIP)– using Louvain-clustering on the cell-topic probability matrix (igraph::cluster_louvain, parameters: k=10, eps=0.1, treetype=“bd”). This led to 55 clusters including the main cell types identified in scRNA-seq. Note that we chose this strategy based on several alternative analyses, in which we observed that cisTopic benefits by higher numbers of cells, even if some of them have few reads, while the clustering of only high-count cells (FIP> 900) provided more stable clusters and more concordant with the scRNA-seq. (2) The same process was then applied to each of the major groups of cells: OL, CB and glia, using separate cisTopic runs, and consensus peaks instead of the ctx pre-defined regions (see section below). These sub-clustering analyses provided 130 clusters, which might be over-clustered, as many of them were not matched to scRNA-seq clusters, but it allowed to identify some extra cell types (e.g. ab-cd split of T4/T5 cells, Extended Data Fig. 6). From these analyses, after the scRNA-seq label transfer (see below), we finally selected 79 clusters as main annotation.

The clusters for the developmental stages were determined following the equivalent approach on the “Larva to 12h APF” cisTopic analysis (in this case only one level of clustering was required).

### Gene accessibility matrix

Gene accessibilities were calculated using the cisTopic probabilities of region accessibility per cell. Next, ctx regions inside the gene body and up to 5kb of its transcription start site were selected. An exponentially decaying function was used to assign distance weights to these regions, were regions further away from the gene have lower weights, similar to ArchR ^104^. To give higher weights to variable regions, we calculated Gini scores per region, where highly variable regions have a high Gini score. Gini scores were then z-standardized and used as exponent for the variability weight. Final weights were defined as the product of the distance and variability weights, and a weighted sum was calculated to acquire a gene accessibility matrix.

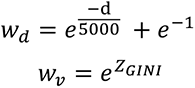

### scRNA clustering

We used the scRNA-data from the whole ageing fly brain from Davie et al.^1^, this time using all data from all protocols with updated analysis methods (mostly batch effect correction). Mapping, filtering, normalization, batch effect correction, clustering, marker gene detection and gene regulatory network inference were performed using the VSN pipeline (https://github.com/vib-singlecell-nf/vsn-pipelines), which is a Nextflow DSL2 pipeline using CellRanger (10x), Scanpy ^105^, Harmony ^106^ and pySCENIC ^45^. We used the command `nextflow -C nextflow.config run vib-singlecell-nf/vsn-pipelines -entry harmony` and a description of the nextflow.config file can be found in Supplementary Data 2. Finally, annotations were transferred from the Davie et al. dataset, by calculating the adjusted rand index between annotations and the different calculated clusterings. The best matching clustering was Leiden resolution 10 (224 clusters) and the clusters were annotated if at least 25% of cells in the cluster had the same annotation. If there was no match, the cluster was retained, ending up with 203 clusters. One modification was made to cluster 15 where a higher resolution (Leiden 12) was chosen in which it split in two (a and b), matching the split detected in the RNA-ATAC co-embedding (see below), leading to the final annotation of 66 clusters out of 204. Subsequently, marker genes were calculated in Seurat’s FindAllMarkers using the Wilcoxon method with min.pct =0.1 and logfc.threshold =0.2).

### Adult cell type annotation: Label transfer from RNA to ATAC (NNLS, AUCell, Seurat)

To assign cell type identities to scATAC-seq clusters (130) we followed three approaches:

First, we used the non-negative least squares method to compare clusters across modalities, similar to what was used in Domcke et al. ^20^. We calculated average RNA expression profiles per cell type from the annotated scRNA-seq data and averaged gene accessibility profiles for the scATAC-seq clusters using the top 10 marker genes per cell type as features (sorted by adjusted p-value (Bonferroni corrected)). These were then used as input for the algorithm in which an optimal weighted sum is calculated and the weights resemble cluster similarities.

Secondly, we used AUCell ^35^ to score gene signatures per cell type based on the top marker genes on the gene accessibility matrix. Gene signatures were then averaged per cluster and clusters were assigned to cell types based on their score.

Thirdly, Seurat v3 ^107^ was used to integrate the gene accessibility and gene expression data. First separate objects were created for scATAC-seq and scRNA-seq data, with the gene accessibility matrix used as “RNA” assay and the region-accessibility as “peaks” assay in the ATAC-seq object. First, the dimensions of the ATAC-seq object were reduced using RunTFIDF, FnidTopFeatures and RunSVD using latent semantic analysis (LSI) on the “peaks” assay with number of components used 50, 70 and 100. Next, the RNA-seq object was log-normalized with NormalizeData with the median of expressed UMIs as scale factor. FindVariableFeatures was used to find 2500 variable features in the RNA-seq data, to be used as features for integration. Anchors for integration were identified using FindTransferAnchors with the RNA-seq data as reference and the ATAC-seq object as query using canonical component analysis using 50, 70 and 100 components. Then TransferData was used to transfer annotations from scRNA-seq to scATAC-seq using the LSI weights for weight reduction and dimensions ranging 2 to 100. To calculate a co-embedding, we used GetAssayData on the variable genes to get the RNA counts and used this as reference data in a second run of TransferData where we impute RNA counts for the ATAC-seq data, again using the LSI weights for weight reduction. The two objects were subsequently merged, followed by scaling of the data, PCA and UMAP and tSNE calculation (ScaleData, RunPCA, RunUMAP, RunTSNE).

Next, we collapsed annotations across all methods and merged non-annotated low-confidence clusters. Tm1/TmY8 and Mi1 matched to the same cluster, but we could separate the sub clusters based on gene accessibility scores of *bsh* and *hth*, two markers for Mi1 neurons.

### omniATAC-seq

Bulk ATAC-seq was performed on 5 samples (three GFP-positive cells from driver lines each targeting one subtype of Kenyon cells (MB371B, MB418B and MB419B) and two negative controls (GFP-negative cells from MB371B and MB419B)). ATAC-seq reads were trimmed using fastq-mcf ^108^ and a list of sequencing primers. The cleaned reads were then used as input for fastqc for quality control. Next, the reads were mapped to the 3rd 2017 FlyBase release (D. melanogaster r6.16) genome using STAR and SAMtools was used to sort the bam file. Macs2 was then used to call differential peaks between the positive samples and their negative controls and both negative controls were used for the positive sample without its own control using `macs2 callpeak -t pos_sample -c neg_sample -g dm –nomodel`.

### Differentially accessible regions and motif analysis

For each of the adult cell clusters (including both clustering resolutions, plus a super-clustering of glia, OL/CB neurons and KCs), we calculated the differentially accessible regions (DARs) based on the predictive distribution from cisTopic (using Wilcoxon rank sum test, run through FindMarkers function in Seurat with the default settings, except logfc.threshold, which was lowered to 0.20, and max.cells.per.ident, which was adjusted to balance the contrasts in some of the analyses). For each of the clusters, the DARs were calculated versus the closest cluster in the tree, and versus all the other clusters in each of the two analyses (i.e. each cluster was compared to rest of the brain, and to the other cells in their same glia/OL/CB/KC category).

We then performed motif enrichment analyses for each of the DAR sets with at least 10 regions using RcisTarget (aucMaxRank=0.01 and 0.05, motif collection version 9, and the TF ChIP-seq database). Each of the analyses was performed comparing versus the whole genome (default settings), and using all the regions in topics as background (re-ranking the database).

The results from these analyses are available in the website (http://flybrain.aertlab.org).

### Cistromes

For building cistromes, we focused on cell types linked to a scRNA-seq cluster, re-grouping the CB clusters into CB-Pros and CB-Imp in order to be able to establish the link to their transcriptome (T4 and T5 cells were analyzed as independent clusters from ATAC, but both mapping to the same T4/T5 RNA cluster).

For each cell type, the cistromes were built based on the motif enrichment analysis of up-regulated DARs sets with at least 10 regions. Each significantly enriched motif (NES>=3) was annotated to expressed TFs based on cisTarget’s “direct” and “inferred by orthology” annotations (considering as expressed TFs those with expression > 0 in at least 10% of the cells of the given type/cluster). Note that since cisTarget’s annotation include some non-TFs DNA binding proteins, we only kept the 459 TFs listed as such on Flybase and GO MF annotation.

For each of those motifs enriched in a DAR-set for a cell type in which the TF is expressed, we retrieved the DARs in which the motif has a significantly high score (i.e. leading edge, see ^102^, using RcisTarget::getSignificantRegions).

The dot-heatmap in Figure 2 shows the average TF expression by cell type (i.e. average of all the cells in the cluster, after normalizing each cell based on its total counts) with max normalization (each gene divided by its maximum value), and the NES of the highest scoring motif (NES capped to 8).

### Gene-enhancer links

We calculated the enhancer-to-gene links using the 43 matched clusters between RNA and ATAC plus CB-Pros and CB-Imp (ATAC T4 and T5 clusters were merged to match T4/T5 in RNA). For each cluster and data modality, 200 pseudocells were created as a bootstrap of 5 cells of the cell type. Each transcriptome pseudocell was then matched to a chromatin pseudocell of the same cell type to calculate the Pearson correlation and Random Forest regression (GENIE3) between each gene’s expression and the predicted accessibility (cisTopic cell-region probability) of the regions within 50kbp of its longest transcript (50kbp upstream the TSS and 50kpb downstream the end, plus the introns). The GENIE3 scores were filtered using the Binarize::binarize.BASC function in R. We then created a score, based on the aggregated ranking of these two measures plus the region accessibility, to allow to select the top regulators. The maximum value of this score was scaled to 1000 for compatibility with UCSC Genome browser (where we suggest a threshold of 600 or 800 for link visualization).

For the comparison of links between BEAF-32 peaks, we used ChIP-seq on whole *Drosophila* embryo (mixed sex embryo of 0-14 hours; ENCODE dataset https://www.encodeproject.org/files/ENCFF704WGH/ ^109^). The peaks were filtered based on the enrichment of the BEAF-32 motif (i-cisTarget analysis with default settings), and their accessibility in the adult fly brain (most of the peaks are ubiquitous across cell types, see Extended Data Fig. 7). We then defined the BEAF-32 based search space for each gene, taking the biggest transcript, and extend (up- and down-stream) until the first BEAF32 peak within 200kbp (skipping the 500bp around the TSS). In case there are no peaks within 200kbp, 50kbp is kept as search space. In 82% of the eGRNs, there is a slightly higher GSEA enrichment score with the TF co-expression module (see below) when using only links within BEAF-32 peaks.

### eGRN integration

The regions in each of the cell-type specific cistromes were converted to genes based on the “enhancer-gene links” with score >= 600. Note that this implies that they may include positive and negative associations. We then used GSEA to check whether each of these gene-sets is enriched in each of the TF co-expression modules from SCENIC-GRNboost (the co-expression modules were used as rankings, after adding the TF itself in the first position if the TF is in any cistrome (i.e. not only on the matching TF)). Therefore, we checked the enrichment of each of the 2110 cell-type specific cistromes with at least 5 target genes (corresponding to 200 TFs) and the 199 TFs also available as co-expression module (using 5000 permutations in GSEA). For each TF co-expression module (i.e. ranking) we kept the significant cistromes for the same TF (p-value < 0.01), and selected the genes in the leading edge to build the e-GRNs. To finalize the e-GRNs per cell type, for each of those genes, we then retrieved the linked regions in the cistrome within the BEAF-32 based search space. Thus, obtaining the connections TF – Region -– Gene.

### eGRN plots

To display the eGRNs as networks in Cytoscape (v3.8.0; ^110^), we focused on the positive region-target gene links (i.e. correlation over 0.22, which corresponds to the top 1% of all calculated gene-region link correlations), and the genes expressed in at least 15% of the cells of the specific cell type (except T4/T5 neurons, for which we used 5% instead).

To reduce the size of the glial networks, they were further filtered by removing TF-region connections with a scaled Cluster-Buster score below 0.25, and regions with a deep learning prediction score below 0.25. The prediction score of each region on the respective deep learning model (Figure 3b; Extended Data Table 2) was calculated with a 500 bp sliding window with a 1 bp shift, and taking the highest score. For each cistrome (i.e. TF and cell type pair), all putative target regions were scored with Cluster-Buster ^111^ using all motifs that had been used to generate the cistrome. For each TF, the highest cluster score per region was kept and scaled between zero and one for the Cytoscape network.

In addition, the Cytoscape networks also display differential expression and accessibility for each node (gene or region, respectively). The differential expression was calculated by contrasting the cell type versus all other cells (avg_logFC calculated with the Seurat function FindMarkers), the accessibility in the cell type was calculated by taking the mean over the interval with subsequent RPGC-normalization.

### Deep learning on topics

Kenyon cells, T-neurons and glia were selected from the adult and 72h APF datasets leading to a total of 17,554 cells. The selected cells were rescored on a set of 207k 150bp peaks (see consensus peaks), which were extended to 300bp for optimal resolution in deep learning. Given the smaller number of cells, we used the conventional Collapsed Gibbs Sampler method in cisTopic using runCGSModels from 1 to 100 topics, with 500 iterations using 250 as burn-in. With selectModel we selected the model with the highest log-likelihood leading to 81 topics. Using runtSNE without PCA on the probability matrix with the cells as target, we acquired the 2D embeddings. We then calculated scores for the topics per region using getRegionsScores with method=’NormTop’ and scale=TRUE. Finally, we used binarizecisTopics with thrP=0.975 to get 81 sets of peaks. These region sets were annotated to the different cell types based on accessibility per cell type and region features (e.g. promotors, BEAF-32) based on motif enrichment and the annotateRegions function using the *Drosophila* datasets.

These sets of regions were then used as input for a deep learning model, where 500bp DNA sequence are used to predict the topic set to which the region belongs. The architecture of the model was used from an earlier study where they again used the cisTopic clusters as an input for the deep learning model (DeepMEL) ^54^. The model is a hybrid CNN-RNN multi-class classifier ^112^, with model architecture details in Extended Data Table 2. In addition to the architecture proposed earlier, we increased the number of filters from 128 to 1024 where 747 of them are initialized as known PWMs representing 212 TFs. Input regions were split into training (80%), validation (10%), and test (10%) sets. The model was trained on the training set, while the validation set was used to do early stopping and to select the best epoch (83rd) as a final model to use.

In order to find the nucleotides that are contributing the most for the topic prediction, we used a network explaining tool called DeepExplainer ^55^. The tool was initialized with 500 random sequences and default parameters were used. The importance score obtained from the DeepExplainer analysis was multiplied by the DNA sequence and visualized as height of the nucleotide letters as in earlier work ^113^. On top of DeepExplainer plots, we performed in-silico saturation mutagenesis where we calculate the effect of each variant of a region on its model prediction score. The sequences with all possible single mutations were generated and delta model score for each topic was calculated.

High nucleotide importances on DeepExplainer plots represent potential binding sites for TFs. We used TF-MoDISco ^56^ to identify the most common patterns for KCs (Topic 21, 35, 77), T-neurons (Topic 23 ,20, 44, 10, 18, 32), and glia (Topic 68, 25, 56, 34, 36). Default parameters were used to run for each group. For conservation study, nucleotides with a DeepExplainer absolute z-standardized importance larger than 3 were selected. The DeepFlyBrain model is deposited in Kipoi (https://kipoi.org/models/DeepFlyBrain).

### Cloning and visualization of enhancers

59 enhancers were chosen based on eGRNs, DARs and development, and the ATAC-seq peak in the targeted region in the cell type of interest was selected. Selected enhancers were scored for the presence of homopolymers (>10) and GC content and small modifications to the sequence were made if needed. Sequences were ordered at Twist Biosciences (US), and inserted in the pTwist ENTR vector. Gateway cloning was then used to insert the sequence in the pH-Stinger vector containing nuclear GFP, Hsp70 promotor and gypsy insulators ^114^. Next, the vectors were sent to FlyORF (CH) and divided into 6 pools that were injected in *Drosophila* embryos. Positive transformants were selected, and PCR was used to determine the identity of the enhancer in each line. This pipeline of pooled injections recovered a transgenic line for 54 of the 59 enhancers. Larval, pupal (24h and 48h) and adult flies were then dissected and stained using the immunohistochemistry protocol for GFP, brp, repo and DAPI. Enhancers were scored using the following system: high off target expression (-2), low off target expression (-1), no expression (0), low on target expression (1) and high on target expression (2). Results can be found in Extended Data Table 4. Tests on the success rate were performed using two-sided Fischer’s exact test.

### Enhancer ROC curves

We used the scikit-learn ^115^ framework to fit a roc-curve to separate adult high quality enhancers (score=2) from the other cloned enhancers. As features we used peak height, peak specificity (Z-score, log-fold change, p-value of DAR), motif content from deep learning, DAR and/or eGRN membership and correlation of peak accessibility with gene expression.

### Annotation of cell types through development

The annotation of cell types through development was performed following two complementary approaches: (1) annotate progenitor cell types based on marker genes near *ase, dpn, grh, dac, cas* and *scro* ^15^ (Extended Data Fig.13 a) and the ventral nerve cord based on abd-A ^15^, and (2) tracking back the annotated adult cell types.

To track back the adult cell types though development we used a Support Vector Machine classifier (SVM): We trained it on the annotated adult cell types, and we used it to iteratively transfer the labels to earlier stages.

1. In the first step, we used the SVM classifier to transfer the labels from the 79 adult clusters (adult cells with more than 900 FIP), to the remaining cells on the adult dataset (adult + 72h APF, the classifiers are trained on the cell-topic matrix). Using cross validation within the adult cells, we estimated that the global accuracy of the classifier is 0.86, with a call rate of 0.97 (it is not forced to assign a class to every cell); having a specificity of over 0.99 for all cell types, and a sensitivity ranging from over 0.90, for many glial, optic lobe and Kenyon cell types, to 0.25-0.50 for the least confident CB-Pros clusters.
2. We then used the adult+72h APF cells to classify the 48h pupa cells using the common cisTopic analysis with these three stages (Extended Data Fig. 2), and Harmony (on the cell-topic matrix) to reduce the effect differences intrinsic to the developmental stage.
3. Finally, we classify the cells in the remaining developmental stages (from larva to 24h AFP). For this we used the global cisTopic analysis (158,116/240,919 cells with more than 900 FIP), with Harmony to correct for developmental stage (Extended Data Fig. 1e). In this last training, we noticed that cells on the progenitor clusters remained largely unassigned, so we finally trained a classifier also including the progenitors as training labels (OL Developing neuron 2, CB Developing neuron 1, OL Neuroepithelium, NB Generation, OL Developing neuron 1, OL Type I NB, CB Type I NB, and LPC), and discarding from the training set the few cells from the “new 48h cluster” that had been assigned to a cell type (they seem to be younger cells, and could distort the classification). This way obtained a likely fate for the developmental cells.

### Core-set identification

The peaks called per cell type per timepoint for the consensus peaks were used as the basis to identify core regions per cell type. Ctx regions (see topic modelling) that overlapped with the called peaks for that timepoint, were defined as open and the DARs of the timepoint for the cell type were used to get differential accessible regions. All ctx regions that passed the filtering were then taken together as one set of total accessible regions of the cell type. Regions that are accessible in every timepoint are defined as the core-set of regions, with regions that are differentially accessible in every timepoint as core-DARs.

### Trajectory of Optic Lobe branches

We used Monocle3 ^116–118^ to fit a trajectory in the 3D UMAP of the optic lobe branch from the larval to 12h APF analysis, and assign pseudotimes to the cells. First, we created a cell_data_set with the region probabilities per cell and the 3D UMAP from cisTopic as embeddings. Then we used cluster_cells followed by partition to separate the optic lobe and central brain branches and subset the object to only contain the optic lobe. We performed another cluster_cell, and selected and merged clusters in the same branch. The branch IDs were then used in Seurat’s FindAllMarkers (Wilcoxon, min.pct=0.1, logfc.threshold=0.25) on the predictive distribution matrix to find differentially accessible regions. Subsequently, motif enrichment on the branch DARs was performed using i- cisTarget ^119^. Regions were linked to genes up to 5kbp up- or downstream and GO was performed using FlyMine (Extended Data Table 5).

### Trajectory of ONE scATAC-seq

The Monocle3 object created for optic lobe branches was also used to calculate pseudotime using learn_graph fit a principal graph and order_cells to assign pseudotimes. Next, optic lobe neuroepithelium cells were selected together with the tips of the lamina precursor cells and optic lobe neuroblasts (NB generation), focusing on the trajectory between these cell types. The trajectory was split into 15 equal parts that were used in Seurat’s FindAllMarkers (Wilcoxon, min.pct=0.1, logfc.threshold=0.1) to find differentially accessible regions with a two-sided Wilcoxon test. Next, the predictive distribution matrix was subset for DARs and CPM normalized, followed by region-based z-normalisation. DARs were grouped into modules using hierarchical clustering with the Scipy cluster.hierachy module ^120^ using distance.pdist (Euclidean), linkage (complete) and fcluster (0.85*max distance) leading to 9 modules. RcisTarget was used to identify motifs per module.

### Trajectory of ONE scRNA-seq

Lamina precursor cells, neuroepithelium cells and optic lobe neuroblasts were selected from a scRNA-seq dataset of the larval brain ^15^. Monocle3 was used to create a trajectory through the cells and assign pseudotimes. First the data was processed using pre_process_cds with principal component analysis as method, selecting 20 components. Next a batch effect correction was performed to align the two different runs with align_cds. The aligned data was then used for reduce_dimension, followed by learn_graph. Once the principal graph was learned, cells were ordered along it and pseudotimes were assigned. To plot gene expression trajectories over pseudotime, a rolling mean was calculated of the log-normalised CPM counts with a window of 10. Next a 10^th^ degree polynomial was fit through the rolling mean with polyfit using NumPy ^121^ and plotted.

### Central brain *pros* vs *Imp*

Central brain clusters in the Adult+72hAPF dataset were selected based on enrichment of central brain only runs (Extended Data Fig. 2), with the exception of Kenyon cells. These clusters were assigned to either *pros* or *Imp* groups based on their maximal mean gene accessibility. We then used Seurat FindAllMarkers on the predictive distribution matrix (Wilcoxon, min.pct=0.1, logfc.threshold=0.2) to identify 166 regions for *pros^+^* cells and 128 regions for *Imp^+^* cells. Motif enrichment was performed using i-cisTarget.

### scATAC-seq embryo

We used scATAC-seq from the whole *Drosophila* embryo ^58^ to map the different central brain cell types. After data download from GEO, we used cisTopic to map the reads on ctx regions leading to 128,510 regions by 20,594 cells matrix. Given the smaller number of cells, we used the conventional Collapsed Gibbs Sampler method in cisTopic using runCGSModels from 1 to 100 topics, with 500 iterations using 250 as burn-in. We selected the model with the highest log-likelihood leading to 50 topics. Using runtSNE without PCA on the probability matrix with the cells as target, we acquired the 2D embeddings. Annotations were transferred from the dataset, identifying the CNS. We then plotted the mean accessibility of the central brain regions on the tSNE.

### Enhancer-switch identification

Region accessibilities per cell type were calculated per timepoint using RPGC normalized bigwig files. Next, a linear curve was fit using statsmodels ^122^ in Python for every region using time as independent variable and region accessibility as dependent variable with 95% confidence intervals calculated for the parameters. Regions with a positive coefficient were assigned to be upregulated and regions with negative coefficients were assigned to be downregulated. Finally, we selected the regions that were upregulated in one cell type while being downregulated in another one, leading to 985 switching regions.

### Cell type specific bams and bigwigs

We used the annotations calculated by the SVM and extracted cells per cell type per timepoint. Next, we subset the bam files from the runs to only contain reads belonging to the selected cells and created a cell type specific bam file. Then we used SAMtools ^123^ to remove duplicates (view -F 0x400) and remove regions mapping blacklisted regions ^124^. This was then used as input for the bamCoverage function from deepTools ^125^ to create an RPGC normalized bigwig file with the following parameters: -`bs 1 -p 8 --normalizeUsing RPGC --effectiveGenomeSize 142573017`.

### Consensus peaks

MACS2 ^126^ was used to call peaks on cell-type specific bam files using the call peak function with following parameters: `macs2 callpeak -q 0.05 -g dm --keep-dup all --nolambda --call- summits --nomodel --shift -75 --extsize 150`. This was repeated for all the timepoints and for the grouped analyses (Adult+P72, L3-P12, P24, P48). Next, the summits were extended to 500bp (or 150bp) using slopBed from BEDTools (-l 149, -r 150 (or -l 74, -r 75). The extended summits were then merged according to ENCODE standards, with first a normalization of the summit score (CPM) followed by iteratively peak merging until non-overlapping peaks across all timepoints and cell types are retained. This led to a final number of 95,921 (500bp) and 207,325 (150bp) peaks.

## Code availability

The updated version of cisTopic for scATAC-seq clustering and topic identification including warpLDA can be found at https://github.com/aertslab/cisTopic with set-up instructions and tutorial. Nextflow pipeline for scRNA-seq analysis can be found at https://github.com/vib-singlecell-nf/vsn-pipelines together with example config files and instructions. The DeepFlyBrain model is deposited in Kipoi (https://kipoi.org/models/DeepFlyBrain). Enhancer gene links can be calculated using ScoMAP (https://github.com/aertslab/ScoMAP) and GENIE3 (https://github.com/aertslab/GENIE3). Trajectory analysis was performed using Monocle3 following the package tutorials (http://cole-trapnell-lab.github.io/monocle-release/monocle3). Differential expression, accessibility and integration of RNA- and ATAC-seq was performed using Seurat v3 (with vignettes and install instructions at https://satijalab.org/seurat/). Code for the website is available at https://github.com/aertslab/FBD_App/.

## Data availability

The data generated for this study have been deposited in NCBI’s Gene Expression Omnibus and are accessible through GEO Series accession number GSE163697. We also provide a dedicated website to browse the results of the analyses and processed data (https://flybrain.aertslab.org), which provides link-outs to the SCope session (http://scope.aertslab.org/#/Fly_Brain/), UCSC hub (http://genome.ucsc.edu/cgi-bin/hgTracks?db=dm6&hubUrl=http://ucsctracks.aertslab.org/papers/FlyBrain/hub.txt), the eGRNs in NDEx, the DeepExplainer plots of enhancers, and other information. The following publicly accessible datasets were also used: GSE107451 (scRNA-seq adult brain), GSE157202 (scRNA-seq larval brain), GSE101581 (scATAC-seq embryo).

## Acknowledgements

This work is funded by the following grants to S. Aerts: ERC Consolidator Grant (724226_cis- CONTROL), by the Special Research Fund (BOF) KU Leuven (grant PF/10/016) and F.W.O (grants G.0791.14, G.0C04.17). J.J. and C.B.G.-B are supported by a PhD fellowship of The Research Foundation – Flanders (FWO, 1199518N and 11F1519N). 10x Chromium was partially made available through VIB Tech Watch Funding. Imaging, FAC-sorting and single-cell analyses were supported by the light microscopy, FACS and single-cell expertise units at the VIB-KU Leuven Center for Brain and Disease Research. Computing was performed at the Vlaams Supercomputer Center (VSC). Stocks obtained from the Bloomington Drosophila Stock Center were used in this study. The funders had no role in study design, data collection and analysis, decision to publish, or preparation of the manuscript. We thank the Janelia FlyLight Project for publicly providing images and reporter lines to assess enhancer activity on the CNS in *Drosophila*. We also thank Filipe Pinto-Teixeira and members of the Aerts lab for helpful discussions, Friday morning éclairs, and for reviewing the manuscript.

## Author contributions

S.Ae., J.J., S.Ai. and D.P. conceived the study. J.J., S.Ai., I.I.T. and D.P. performed computational analyses with assistance from K.S., C.B.G.-B, G.H. and M.D. S.M. and V.C. performed scATAC-seq experiments, J.J. and S.M. performed FAC-sorting and omniATAC-seq. J.N.I., J.J, X.J.Q., and S.M. performed antibody staining and visualization. X.J.Q. and V.C. performed the cloning of selected enhancers. S.Ai. created the website with assistance of G.H., D.P. and K.S. S.Ae., J.J., S.Ai. and I.I.T. wrote the manuscript.

## List of Extended Data Tables

**Extended Data Table 1**: CellRanger statistics of the 10x Chromium runs.

**Extended Data Table 2**: Architecture and hyperparameters of DeepFlyBrain.

**Extended Data Table 3**: Topic annotations and performance metrics of DeepFlyBrain.

**Extended Data Table 4**: Cloned enhancers and transgenic lines made for in vivo reporter assays.

**Extended Data Table 5**: GO results for optic lobe branches.

## List of supplementary files

**Supplementary Data 1**: FACS Gating Strategy

**Supplementary Data 2**: VSN config file

**Supplementary Data 3**: DeepFlyBrain training

**Supplementary Data 4**: DeepFlyBrain performance

**Supplementary Data 5**: DeepFlyBrain scoring and DeepExplainer plots

## References

1. Davie, K. et al. A Single-Cell Transcriptome Atlas of the Aging Drosophila Brain. Cell 174, 982–998.e20 (2018).

2. Konstantinides, N. et al. Phenotypic Convergence: Distinct Transcription Factors Regulate Common Terminal Features. Cell 174, 622–635.e13 (2018).

3. Croset, V., Treiber, C. D. & Waddell, S. Cellular diversity in the Drosophila midbrain revealed by single-cell transcriptomics. eLife 7, e34550 (2018).

4. Özel, M. N. et al. Neuronal diversity and convergence in a visual system developmental atlas. Nature 1–8 (2020) doi:10.1038/s41586-020-2879-3.

5. Kurmangaliyev, Y. Z., Yoo, J., Valdes-Aleman, J., Sanfilippo, P. & Zipursky, S. L. Transcriptional Programs of Circuit Assembly in the Drosophila Visual System. Neuron (2020) doi:10.1016/j.neuron.2020.10.006.

6. Costa, M., Manton, J. D., Ostrovsky, A. D., Prohaska, S. & Jefferis, G. S. X. E. NBLAST: Rapid, Sensitive Comparison of Neuronal Structure and Construction of Neuron Family Databases. Neuron 91, 293–311 (2016).

7. Rivera-Alba, M. et al. Wiring economy and volume exclusion determine neuronal placement in the Drosophila brain. Curr. Biol. 21, 2000–2005 (2011).

8. Scheffer, L. K. et al. A connectome and analysis of the adult Drosophila central brain. eLife 9, (2020).

9. Xu, C. S. et al. A Connectome of the Adult Drosophila Central Brain. bioRxiv 2020.01.21.911859 (2020) doi:10.1101/2020.01.21.911859.

10. Zheng, Z. et al. A Complete Electron Microscopy Volume of the Brain of Adult Drosophila melanogaster. Cell 174, 730–743.e22 (2018).

11. Jenett, A. et al. A GAL4-Driver Line Resource for Drosophila Neurobiology. Cell Rep 2, 991–1001 (2012).

12. Robie, A. A. et al. Mapping the Neural Substrates of Behavior. Cell 170, 393–406.e28 (2017).

13. Brunet Avalos, C., Maier, G. L., Bruggmann, R. & Sprecher, S. G. Single cell transcriptome atlas of the Drosophila larval brain. Elife 8, (2019).

14. Cocanougher, B. T. et al. Comparative single-cell transcriptomics of complete insect nervous systems. bioRxiv 785931 (2020) doi:10.1101/785931.

15. Ravenscroft, T. A. et al. Drosophila Voltage-Gated Sodium Channels Are Only Expressed in Active Neurons and Are Localized to Distal Axonal Initial Segment-like Domains. J. Neurosci. 40, 7999– 8024 (2020).

16. Allen, A. M. et al. A single-cell transcriptomic atlas of the adult Drosophila ventral nerve cord. eLife 9, e54074 (2020).

17. Buenrostro, J. D., Giresi, P. G., Zaba, L. C., Chang, H. Y. & Greenleaf, W. J. Transposition of native chromatin for fast and sensitive epigenomic profiling of open chromatin, DNA-binding proteins and nucleosome position. Nat. Methods 10, 1213–1218 (2013).

18. Buenrostro, J. D. et al. Single-cell chromatin accessibility reveals principles of regulatory variation. Nature 523, 486–490 (2015).

19. Cao, J. et al. A human cell atlas of fetal gene expression. Science 370, (2020).

20. Domcke, S. et al. A human cell atlas of fetal chromatin accessibility. Science 370, (2020).

21. Lake, B. B. et al. Integrative single-cell analysis of transcriptional and epigenetic states in the human adult brain. Nat. Biotechnol. 36, 70–80 (2018).

22. Doe, C. Q. Temporal Patterning in the Drosophila CNS. Annu. Rev. Cell Dev. Biol. 33, 219–240 (2017).

23. Erclik, T. et al. Integration of temporal and spatial patterning generates neural diversity. Nature 541, 365–370 (2017).

24. Estacio-Gómez, A., Hassan, A., Walmsley, E., Le, L. W. & Southall, T. D. Dynamic neurotransmitter specific transcription factor expression profiles during Drosophila development. Biology Open 9, (2020).

25. Komiyama, T., Johnson, W. A., Luo, L. & Jefferis, G. S. X. E. From lineage to wiring specificity. POU domain transcription factors control precise connections of Drosophila olfactory projection neurons. Cell 112, 157–167 (2003).

26. Kurmangaliyev, Y. Z., Yoo, J., LoCascio, S. A. & Zipursky, S. L. Modular transcriptional programs separately define axon and dendrite connectivity. Elife 8, (2019).

27. Schilling, T., Ali, A. H., Leonhardt, A., Borst, A. & Pujol-Martí, J. Transcriptional control of morphological properties of direction-selective T4/T5 neurons in Drosophila. Development 146, (2019).

28. Halder, G., Callaerts, P. & Gehring, W. J. Induction of ectopic eyes by targeted expression of the eyeless gene in Drosophila. Science 267, 1788–1792 (1995).

29. Masserdotti, G., Gascón, S. & Götz, M. Direct neuronal reprogramming: learning from and for development. Development 143, 2494–2510 (2016).

30. Hecker, M., Lambeck, S., Toepfer, S., van Someren, E. & Guthke, R. Gene regulatory network inference: Data integration in dynamic models—A review. Biosystems 96, 86–103 (2009).

31. Li, Z. et al. Identification of transcription factor binding sites using ATAC-seq. Genome Biology 20, 45 (2019).

32. Li, H. et al. Classifying Drosophila Olfactory Projection Neuron Subtypes by Single-Cell RNA Sequencing. Cell 171, 1206–1220.e22 (2017).

33. Mackay, T. F. C. et al. The Drosophila melanogaster Genetic Reference Panel. Nature 482, 173– 178 (2012).

34. Bravo González-Blas, C., et al. cisTopic: cis-regulatory topic modeling on single-cell ATAC-seq data. Nature Methods 16, 397–400 (2019).

35. Aibar, S. et al. SCENIC: single-cell regulatory network inference and clustering. Nature Methods 14, 1083–1086 (2017).

36. Stanescu, D. E., Yu, R., Won, K.-J. & Stoffers, D. A. Single cell transcriptomic profiling of mouse pancreatic progenitors. Physiol Genomics 49, 105–114 (2017).

37. Shih, M.-F. M., Davis, F. P., Henry, G. L. & Dubnau, J. Nuclear Transcriptomes of the Seven Neuronal Cell Types That Constitute the Drosophila Mushroom Bodies. G3 (Bethesda) 9, 81–94 (2019).

38. Corces, M. R. et al. An improved ATAC-seq protocol reduces background and enables interrogation of frozen tissues. Nature Methods 14, 959–962 (2017).

39. Trevino, A. E. et al. Chromatin accessibility dynamics in a model of human forebrain development. Science 367, (2020).

40. Davis, F. P. et al. A genetic, genomic, and computational resource for exploring neural circuit function. eLife 9, e50901 (2020).

41. Huynh-Thu, V. A., Irrthum, A., Wehenkel, L. & Geurts, P. Inferring Regulatory Networks from Expression Data Using Tree-Based Methods. PLOS ONE 5, e12776 (2010).

42. Iacono, G., Massoni-Badosa, R. & Heyn, H. Single-cell transcriptomics unveils gene regulatory network plasticity. Genome Biology 20, 110 (2019).

43. Jackson, C. A., Castro, D. M., Saldi, G.-A., Bonneau, R. & Gresham, D. Gene regulatory network reconstruction using single-cell RNA sequencing of barcoded genotypes in diverse environments. eLife 9, e51254 (2020).

44. Matsumoto, H. et al. SCODE: an efficient regulatory network inference algorithm from single-cell RNA-Seq during differentiation. Bioinformatics 33, 2314–2321 (2017).

45. Van de Sande, B. et al. A scalable SCENIC workflow for single-cell gene regulatory network analysis. Nature Protocols 15, 2247–2276 (2020).

46. Crittenden, J. R., Skoulakis, E. M. C., Goldstein, E. S. & Davis, R. L. Drosophila mef2 is essential for normal mushroom body and wing development. Biology Open 7, (2018).

47. Schulz, R. A., Chromey, C., Lu, M. F., Zhao, B. & Olson, E. N. Expression of the D-MEF2 transcription in the Drosophila brain suggests a role in neuronal cell differentiation. Oncogene 12, 1827–1831 (1996).

48. Minocha, S., Boll, W. & Noll, M. Crucial roles of Pox neuro in the developing ellipsoid body and antennal lobes of the Drosophila brain. PLoS One 12, (2017).

49. Naidu, V. G. et al. Temporal progression of Drosophila medulla neuroblasts generates the transcription factor combination to control T1 neuron morphogenesis. Developmental Biology 464, 35–44 (2020).

50. Avet-Rochex, A., Maierbrugger, K. T. & Bateman, J. M. Glial enriched gene expression profiling identifies novel factors regulating the proliferation of specific glial subtypes in the Drosophila brain. Gene Expr Patterns 16, 61–68 (2014).

51. Harmston, N. et al. Topologically associating domains are ancient features that coincide with Metazoan clusters of extreme noncoding conservation. Nature Communications 8, 441 (2017).

52. Wang, Q., Sun, Q., Czajkowsky, D. M. & Shao, Z. Sub-kb Hi-C in D . melanogaster reveals conserved characteristics of TADs between insect and mammalian cells. Nature Communications 9, 188 (2018).

53. Yang, J., Ramos, E. & Corces, V. G. The BEAF-32 insulator coordinates genome organization and function during the evolution of Drosophila species. Genome Res 22, 2199–2207 (2012).

54. Minnoye, L. et al. Cross-species analysis of enhancer logic using deep learning. Genome Res. gr.260844.120 (2020) doi:10.1101/gr.260844.120.

55. Lundberg, S. M. & Lee, S.-I. A Unified Approach to Interpreting Model Predictions. Advances in Neural Information Processing Systems 30, 4765–4774 (2017).

56. Shrikumar, A. et al. Technical Note on Transcription Factor Motif Discovery from Importance Scores (TF-MoDISco) version 0.5.6.5. arXiv:1811.00416 [cs, q-bio, stat] (2020).

57. Hubisz, M. J., Pollard, K. S. & Siepel, A. PHAST and RPHAST: phylogenetic analysis with space/time models. Brief Bioinform 12, 41–51 (2011).

58. Cusanovich, D. A. et al. The cis -regulatory dynamics of embryonic development at single-cell resolution. Nature 555, 538–542 (2018).

59. Bravo González-Blas, C., et al. Identification of genomic enhancers through spatial integration of single-cell transcriptomics and epigenomics. Molecular Systems Biology 16, e9438 (2020).

60. Apitz, H. & Salecker, I. A challenge of numbers and diversity: neurogenesis in the Drosophila optic lobe. J Neurogenet 28, 233–249 (2014).

61. Chotard, C., Leung, W. & Salecker, I. glial cells missing and gcm2 Cell Autonomously Regulate Both Glial and Neuronal Development in the Visual System of Drosophila. Neuron 48, 237–251 (2005).

62. Endo, K. et al. Chromatin modification of Notch targets in olfactory receptor neuron diversification. Nature Neuroscience 15, 224–233 (2012).

63. Eroglu, E. et al. SWI/SNF Complex Prevents Lineage Reversion and Induces Temporal Patterning in Neural Stem Cells. Cell 156, 1259–1273 (2014).

64. Piñeiro, C., Lopes, C. S. & Casares, F. A conserved transcriptional network regulates lamina development in the Drosophila visual system. Development 141, 2838–2847 (2014).

65. Lee, J., Park, S.-Y. & Yoo, S. Roles of Nk2.1/scro homeobox gene in the development of optic lobe neuroblast in Drosophila melanogaster. IBRO Reports 6, S339–S340 (2019).

66. Yoo, S. et al. Knock-in mutations of scarecrow, a Drosophila homolog of mammalian Nkx2.1, reveal a novel function required for development of the optic lobe in Drosophila melanogaster. Developmental Biology 461, 145–159 (2020).

67. Medioni, C., Ramialison, M., Ephrussi, A. & Besse, F. Imp Promotes Axonal Remodeling by Regulating profilin mRNA during Brain Development. Current Biology 24, 793–800 (2014).

68. Vijayakumar, J. et al. The prion-like domain of Drosophila Imp promotes axonal transport of RNP granules in vivo. Nature Communications 10, 2593 (2019).

69. Alyagor, I. et al. Combining Developmental and Perturbation-Seq Uncovers Transcriptional Modules Orchestrating Neuronal Remodeling. Developmental Cell 47, 38–52.e6 (2018).

70. Kirilly, D. et al. A genetic pathway composed of Sox14 and Mical governs severing of dendrites during pruning. Nat. Neurosci. 12, 1497–1505 (2009).

71. Cheng, S. et al. Molecular basis of synaptic specificity by immunoglobulin superfamily receptors in Drosophila. eLife 8, e41028 (2019).

72. Tan, L. et al. Ig Superfamily Ligand and Receptor Pairs Expressed in Synaptic Partners in Drosophila. Cell 163, 1756–1769 (2015).

73. Ngo, K. T., Andrade, I. & Hartenstein, V. Spatio-temporal pattern of neuronal differentiation in the Drosophila visual system: A user’s guide to the dynamic morphology of the developing optic lobe. Developmental Biology 428, 1–24 (2017).

74. Ma, S. et al. Chromatin Potential Identified by Shared Single-Cell Profiling of RNA and Chromatin. Cell 183, 1103–1116.e20 (2020).

75. Cusanovich, D. A. et al. A Single-Cell Atlas of In Vivo Mammalian Chromatin Accessibility. Cell 174, 1309–1324.e18 (2018).

76. Preissl, S. et al. Single-nucleus analysis of accessible chromatin in developing mouse forebrain reveals cell-type-specific transcriptional regulation. Nat Neurosci 21, 432–439 (2018).

77. Chen, S., Lake, B. B. & Zhang, K. High-throughput sequencing of the transcriptome and chromatin accessibility in the same cell. Nature Biotechnology 37, 1452–1457 (2019).

78. Zhu, C. et al. An ultra high-throughput method for single-cell joint analysis of open chromatin and transcriptome. Nature Structural & Molecular Biology 26, 1063–1070 (2019).

79. Berg, O. G. & von Hippel, P. H. Selection of DNA binding sites by regulatory proteins. Statistical-mechanical theory and application to operators and promoters. J Mol Biol 193, 723–750 (1987).

80. Crocker, J., Preger-Ben Noon, E. & Stern, D. L. Chapter Twenty-Seven - The Soft Touch: Low-Affinity Transcription Factor Binding Sites in Development and Evolution. in Current Topics in Developmental Biology (ed. Wassarman, P. M.) vol. 117 455–469 (Academic Press, 2016).

81. Kribelbauer, J. F., Rastogi, C., Bussemaker, H. J. & Mann, R. S. Low-Affinity Binding Sites and the Transcription Factor Specificity Paradox in Eukaryotes. Annu Rev Cell Dev Biol 35, 357–379 (2019).

82. Scardigli, R., Bäumer, N., Gruss, P., Guillemot, F. & Le Roux, I. Direct and concentration-dependent regulation of the proneural gene Neurogenin2 by Pax6. Development 130, 3269– 3281 (2003).

83. Koromila, T. et al. Odd-paired is a pioneer-like factor that coordinates with Zelda to control gene expression in embryos. eLife 9, e59610 (2020).

84. Kudron, M. M. et al. The ModERN Resource: Genome-Wide Binding Profiles for Hundreds of Drosophila and Caenorhabditis elegans Transcription Factors. Genetics 208, 937–949 (2018).

85. Ozdemir, A., Ma, L., White, K. P. & Stathopoulos, A. Su(H)-mediated repression positions gene boundaries along the dorsal-ventral axis of Drosophila embryos. Dev Cell 31, 100–113 (2014).

86. Samata, M. et al. Intergenerationally Maintained Histone H4 Lysine 16 Acetylation Is Instructive for Future Gene Activation. Cell 182, 127–144.e23 (2020).

87. Ye, Y. et al. Chromatin remodeling during the in vivo glial differentiation in early Drosophila embryos. Scientific Reports 6, 33422 (2016).

88. Brás-Pereira, C. et al. dachshund Potentiates Hedgehog Signaling during Drosophila Retinogenesis. PLoS Genet 12, (2016).

89. Dardalhon-Cuménal, D. et al. Cyclin G and the Polycomb Repressive complexes PRC1 and PR-DUB cooperate for developmental stability. PLOS Genetics 14, e1007498 (2018).

90. Donohoe, C. D. et al. Atf3 links loss of epithelial polarity to defects in cell differentiation and cytoarchitecture. PLoS Genet 14, (2018).

91. Jusiak, B. et al. Regulation of Drosophila Eye Development by the Transcription Factor Sine oculis. PLOS ONE 9, e89695 (2014).

92. Newcomb, S. et al. cis-regulatory architecture of a short-range EGFR organizing center in the Drosophila melanogaster leg. PLoS Genet 14, e1007568 (2018).

93. Schertel, C. et al. A large-scale, in vivo transcription factor screen defines bivalent chromatin as a key property of regulatory factors mediating Drosophila wing development. Genome Res 25, 514–523 (2015).

94. Yeung, K. et al. Integrative genomic analysis reveals novel regulatory mechanisms of eyeless during Drosophila eye development. Nucleic Acids Res 46, 11743–11758 (2018).

95. Koemans, T. S. et al. Functional convergence of histone methyltransferases EHMT1 and KMT2C involved in intellectual disability and autism spectrum disorder. PLoS Genet 13, e1006864 (2017).

96. Magadi, S. S. et al. Dissecting Hes-centred transcriptional networks in neural stem cell maintenance and tumorigenesis in Drosophila. Development 147, (2020).

97. Ray, P. et al. Combgap contributes to recruitment of Polycomb group proteins in Drosophila. Proc Natl Acad Sci U S A 113, 3826–3831 (2016).

98. Brand, A. H. & Perrimon, N. Targeted gene expression as a means of altering cell fates and generating dominant phenotypes. Development 118, 401–415 (1993).

99. Boll, W. & Noll, M. The Drosophila Pox neuro gene: control of male courtship behavior and fertility as revealed by a complete dissection of all enhancers. Development 129, 5667–5681 (2002).

100. Gramates, L. S. et al. FlyBase at 25: looking to the future. Nucleic Acids Res 45, D663–D671 (2017).

101. Kang, H. M. et al. Multiplexed droplet single-cell RNA-sequencing using natural genetic variation. Nature Biotechnology 36, 89–94 (2018).

102. Herrmann, C., Van de Sande, B., Potier, D. & Aerts, S. i-cisTarget: an integrative genomics method for the prediction of regulatory features and cis-regulatory modules. Nucleic Acids Res 40, e114 (2012).

103. Chen, J., Li, K., Zhu, J. & Chen, W. WarpLDA: a cache efficient O(1) algorithm for latent dirichlet allocation. Proc. VLDB Endow. 9, 744–755 (2016).

104. Granja, J. M. et al. ArchR: An integrative and scalable software package for single-cell chromatin accessibility analysis. bioRxiv 2020.04.28.066498 (2020) doi:10.1101/2020.04.28.066498.

105. Wolf, F. A., Angerer, P. & Theis, F. J. SCANPY: large-scale single-cell gene expression data analysis. Genome Biology 19, 15 (2018).

106. Korsunsky, I. et al. Fast, sensitive and accurate integration of single-cell data with Harmony. Nature Methods 16, 1289–1296 (2019).

107. Stuart, T. et al. Comprehensive Integration of Single-Cell Data. Cell 177, 1888–1902.e21 (2019).

108. Aronesty et al. ea-utils: ‘Command-line tools for processing biological sequencing data’. (2011).

109. Davis, C. A. et al. The Encyclopedia of DNA elements (ENCODE): data portal update. Nucleic Acids Res 46, D794–D801 (2018).

110. Shannon, P. et al. Cytoscape: a software environment for integrated models of biomolecular interaction networks. Genome Res 13, 2498–2504 (2003).

111. Frith, M. C., Li, M. C. & Weng, Z. Cluster-Buster: Finding dense clusters of motifs in DNA sequences. Nucleic Acids Res 31, 3666–3668 (2003).

112. Quang, D. & Xie, X. DanQ: a hybrid convolutional and recurrent deep neural network for quantifying the function of DNA sequences. Nucleic Acids Res 44, e107 (2016).

113. Shrikumar, A., Greenside, P. & Kundaje, A. Learning Important Features Through Propagating Activation Differences. arXiv:1704.02685 [cs] (2019).

114. Aerts, S. et al. Robust Target Gene Discovery through Transcriptome Perturbations and Genome-Wide Enhancer Predictions in Drosophila Uncovers a Regulatory Basis for Sensory Specification. PLOS Biology 8, e1000435 (2010).

115. Pedregosa, F. et al. Scikit-learn: Machine Learning in Python. Journal of Machine Learning Research 12, 2825–2830 (2011).

116. Cao, J. et al. The single-cell transcriptional landscape of mammalian organogenesis. Nature 566, 496–502 (2019).

117. Qiu, X. et al. Reversed graph embedding resolves complex single-cell trajectories. Nature Methods 14, 979–982 (2017).

118. Trapnell, C. et al. The dynamics and regulators of cell fate decisions are revealed by pseudotemporal ordering of single cells. Nature Biotechnology 32, 381–386 (2014).

119. Imrichová, H., Hulselmans, G., Kalender Atak, Z., Potier, D. & Aerts, S. i-cisTarget 2015 update: generalized cis-regulatory enrichment analysis in human, mouse and fly. Nucleic Acids Res 43, W57–W64 (2015).

120. Virtanen, P. et al. SciPy 1.0: fundamental algorithms for scientific computing in Python. Nature Methods 17, 261–272 (2020).

121. Harris, C. R. et al. Array programming with NumPy. Nature 585, 357–362 (2020).

122. Seabold, S. & Perktold, J. Statsmodels: Econometric and Statistical Modeling with Python. in 92–96 (2010). doi:10.25080/Majora-92bf1922-011.

123. Li, H. et al. The Sequence Alignment/Map format and SAMtools. Bioinformatics 25, 2078–2079 (2009).

124. Amemiya, H. M., Kundaje, A. & Boyle, A. P. The ENCODE Blacklist: Identification of Problematic Regions of the Genome. Scientific Reports 9, 9354 (2019).

125. Ramírez, F. et al. deepTools2: a next generation web server for deep-sequencing data analysis. Nucleic Acids Research 44, W160–W165 (2016).

126. Zhang, Y. et al. Model-based Analysis of ChIP-Seq (MACS). Genome Biology 9, R137 (2008).

